# Notch inhibition promotes regeneration and immunosuppression supports cone survival in a zebrafish model of inherited retinal dystrophy

**DOI:** 10.1101/2021.12.19.473212

**Authors:** Joseph Fogerty, Ping Song, Patrick Boyd, Sarah Grabinski, Thanh Hoang, Adrian Reich, Lauren T. Cianciolo, Seth Blackshaw, Jeff S. Mumm, David R. Hyde, Brian D. Perkins

## Abstract

Photoreceptor degeneration leads to irreversible vision loss in humans with retinal dystrophies such as Retinitis Pigmentosa. Whereas photoreceptor loss is permanent in mammals, zebrafish possesses the ability to regenerate retinal neurons and restore visual function. Following acute damage, Müller glia (MG) re-enter the cell cycle and produce multipotent progenitors whose progeny differentiate into mature neurons. Both MG reprogramming and proliferation of retinal progenitor cells require reactive microglia and associated inflammatory signaling. Paradoxically, MG in zebrafish models of photoreceptor degeneration fail to re-enter the cell cycle and regenerate lost cells. Here, we used the zebrafish *cep290* mutant to demonstrate that progressive cone degeneration generates an immune response but does not stimulate MG proliferation. Acute light damage triggered photoreceptor regeneration in *cep290* mutants but cones were only restored to pre-lesion densities. Using *irf8* mutant zebrafish, we found that the chronic absence of microglia reduced inflammation and rescued cone degeneration in *cep290* mutants. Finally, single-cell RNA-sequencing revealed sustained expression of *notch3* in MG of *cep290* mutants and inhibition of Notch signaling induced MG to re-enter the cell cycle. Our findings provide new insights on the requirements for MG to proliferate and the potential for immunosuppression to prolong photoreceptor survival.

## Introduction

In humans and mammals, the loss of retinal neurons from injury or inherited retinal degeneration results in visual impairment or blindness. Vision loss is irreversible because mammals do not mount a regenerative response to lost or dying neurons. In contrast, fish possess the capability to regenerate lost neurons and restore visual function from populations of endogenous stem cells (1–10). Numerous groups have revealed the identity of retina stem cell populations (11, 12), identified the sequence of events that underpin regeneration (13, 14), and dissected several molecular pathways that promote and inhibit regeneration (15–19).

In zebrafish, Müller glia (MG) constitute an inducible stem cell population capable of regenerating lost neurons. In the uninjured retina, MG periodically divide asymmetrically to produce a rod-committed progenitor cell (20–22). These cells migrate to the outer nuclear layer (ONL), whereupon they are referred to as rod precursors (23). These rod precursors may continue to proliferate or undergo a terminal mitosis and differentiate into rod photoreceptors (10, 20, 22). In response to widespread acute retinal damage, however, MG undergo cellular reprogramming and produce multipotent retinal progenitors that proliferate and differentiate into all retinal cell types (2, 24). If damage is limited to a small number of rod photoreceptors, proliferation of rod precursors will increase without a noticeable increase in MG proliferation (25–27). Conversely, cone photoreceptors can only be regenerated from MG-derived retinal progenitors in the adult retina.

The contribution of immune cells and inflammation to both degeneration and regeneration has garnered considerable interest in recent years. In response to photoreceptor degeneration in mammals, microglia, the resident immune cell of the retina, become activated, release pro-inflammatory cytokines, and phagocytize photoreceptors (28–30). In addition, recent in vivo imaging studies revealed that peripheral monocytes also infiltrate the mouse retina early in the degeneration process (31). Zebrafish also exhibit a robust inflammatory response following acute retinal injury, with activation of microglia/macrophages and the release of pro-inflammatory cytokines (13, 32–34). Perhaps because the zebrafish retina is avascular, this immune response is driven primarily by resident microglia (32). However, mechanical or chemical damage that compromises the blood-retinal-barrier can allow circulating monocytes, macrophages, and neutrophils to infiltrate the retina and contribute to inflammation (33). While microglia-mediated inflammation is generally considered neurotoxic in mammals, inflammation in zebrafish appears critical for the reprogramming of MG and proliferation of MG-derived progenitors (13, 35). Pharmacological immunosuppression with dexamethasone or depletion of microglia with PLX3397 significantly impaired proliferation of MG-derived progenitors, thereby restricting regeneration (32, 35, 36). As immunomodulation has been posited as a potential therapy for retinal degeneration (37, 38), it is important to consider the impact of immunosuppression on regeneration in zebrafish models of progressive retinal degeneration.

Although it is typically assumed that zebrafish regenerate retinal neurons following damage and disease, knowledge about regeneration comes primarily from studies where the retina was acutely damaged by light, mechanical injury, or injection of toxins such as ouabain (5, 9, 16, 24). Zebrafish harboring mutations in genes associated with inherited retinal degeneration frequently die during larval stages before robust regeneration occurs (39–48). Multiple reports exist, however, of adult zebrafish models with progressive photoreceptor degeneration with little evidence for cone regeneration from MG-derived progenitors (49–56). In the *gold rush* (*gosh*; *aryl hydrocarbon receptor interacting protein like 2*) and *bbs2* (*Bardet-Biedl Syndrome 2*) mutants, proliferation of MG could be induced by injection of exogenous TNFα or light damage, respectively (55, 56), indicating that the ability of MG to proliferate persists. Here, we utilize a zebrafish *cep290* mutant (*centrosomal protein 290*) to determine the immune response in a model of progressive retinal degeneration. We demonstrate that progressive retinal degeneration triggers a transcriptional and cellular immune response. We show that inflammation alone is insufficient to trigger MG reprogramming. Whereas immunosuppression by dexamethasone inhibited proliferation of rod precursor cells, blocking glucocorticoid signaling with RU486 did not stimulate regeneration. Loss of *irf8* in addition to *cep290* resulted in chronic immunosuppression and prevented cone degeneration. Finally, we show that upregulation of *notch3* in MG of *cep290* mutants suppresses their proliferation, and that Notch inhibition releases that constraint. Collectively, our results indicate that therapies to suppress microglia function may prolong photoreceptor survival and that triggering regeneration requires more than inflammation alone.

## Results

### Proliferating progenitor cells differentiate into rod photoreceptors in *cep290* mutants

We previously reported that the *cep290* mutant undergoes progressive cone degeneration and noted an increased number of cells in the ONL that stained positive for the proliferation marker PCNA (57). This suggested that rod photoreceptors also degenerated but were replaced by unipotent rod precursors (10, 22) and that degeneration of photoreceptors remained insufficient to induce MG proliferation in *cep290* mutants. To investigate the identity of the proliferating cells and to confirm that MG were not undergoing mitosis, the *cep290* mutant was crossed into the transgenic reporter line *Tg(gfap:eGFP)mi2002*, which labels MG with GFP (58). At 6 months post fertilization (mpf), *cep290* mutants and wild-type siblings were injected with EdU and eyes were collected for immunohistochemistry 24 hours later. In wild-type retinas, a few EdU+ cells were present in the ONL and inner nuclear layer (INL) (SI Appendix, Fig. S1A). In *cep290* mutants, considerably more EdU+ cells were observed in the ONL (SI Appendix, Fig. S1B). None of the EdU+ cells colocalized with GFP+ MG nuclei in the INL. These results strongly suggest that proliferation is limited to ONL-localized rod precursors in *cep290* mutants. We next sought to confirm that the proliferating cells gave rise exclusively to rod photoreceptors. To unambiguously label rod nuclei, the *cep290* mutation was crossed with the transgenic reporter line *Tg(Xla.Rho:eGFP)fl1*, also known as *Tg(XOPS:eGFP)* (59), and animals were injected with EdU. Retinal cryosections were processed and imaged for EdU, GFP fluorescence (rods), and with the antibody Zpr-1 (red/green double cone photoreceptors). EdU colocalized with GFP from rod photoreceptors (SI Appendix, Fig. S1C) and failed to localize to Zpr-1+ cones (SI Appendix, Fig. S1D). The GFP fluorescence intensity was measured in individual EdU+ cells, in cells located in the INL, and in Zpr-1+ cells and normalized to a sample of GFP+ cells in the ONL from the same image (SI Appendix, Fig. S1E). Cells in the INL and all Zpr-1+ cells were EdU-, whereas all EdU+ cells exhibited GFP signal above background. These data indicate that proliferation of rod precursors in *cep290* mutants contributes to regeneration of rods and likely explains why rod degeneration is not observed in *cep290* mutants.

### Progressive retinal degeneration in *cep290* mutants leads to an immune cell response

Retinal injury and cell death in zebrafish results in rapid activation of resident microglia and increased expression of inflammatory cytokines, such as TNFα (13, 32, 33). Activation of microglia is associated with proliferation of MG and techniques that ablate or deplete microglia attenuate retinal regeneration following damage (32, 36). To determine if *cep290* mutants exhibited signs of inflammation, retinas from 6 mpf animals were stained with the antibody 4C4 to label microglia/macrophages (60), and anti-L-plastin antibodies to label leukocytes (33, 61). In wild-type retinas, ramified microglia/macrophage were found in the nerve fiber layer, the inner plexiform layer (IPL) and outer plexiform layer (OPL), and in the subretinal space between the outer segments and RPE (Fig. 1A). In contrast, significant numbers of amoeboid-shaped, activated microglia/macrophage accumulated in the subretinal space of *cep290* mutants (Fig. 1B). These cells were both 4C4^+^ and L-plastin^+^, which is consistent with resident microglia/macrophages responding to photoreceptor degeneration. Similar to prior results from rod ablation studies in larvae (32), no evidence was found that peripheral macrophages (e.g. 4C4^-^,L-plastin^+^ cells) entered the retinas of *cep290* mutants. In both wild-type and *cep290* mutants, microglia/macrophages in the OPL remained ramified (Figs. 1C and D), although an 80% reduction in microglia/macrophages was observed in the IPL of *cep290* mutants (cyan arrows, Figs. 1C, 1D; quantified Fig. 1E). This resembles what occurs in humans with retinitis pigmentosa and in rodent models of inherited retinal degeneration, where death of photoreceptors promotes the translocation of activated microglia/macrophages from the inner retina to the subretinal space and the release of inflammatory cytokines, including TNFα (38, 62). Combined with the results above, these data indicate that although *cep290* mutants exhibited immune system activation and proliferation of rod precursors, the level of immune reactivity was insufficient to induce the mutant MG to enter the cell cycle.

**Figure 1.**
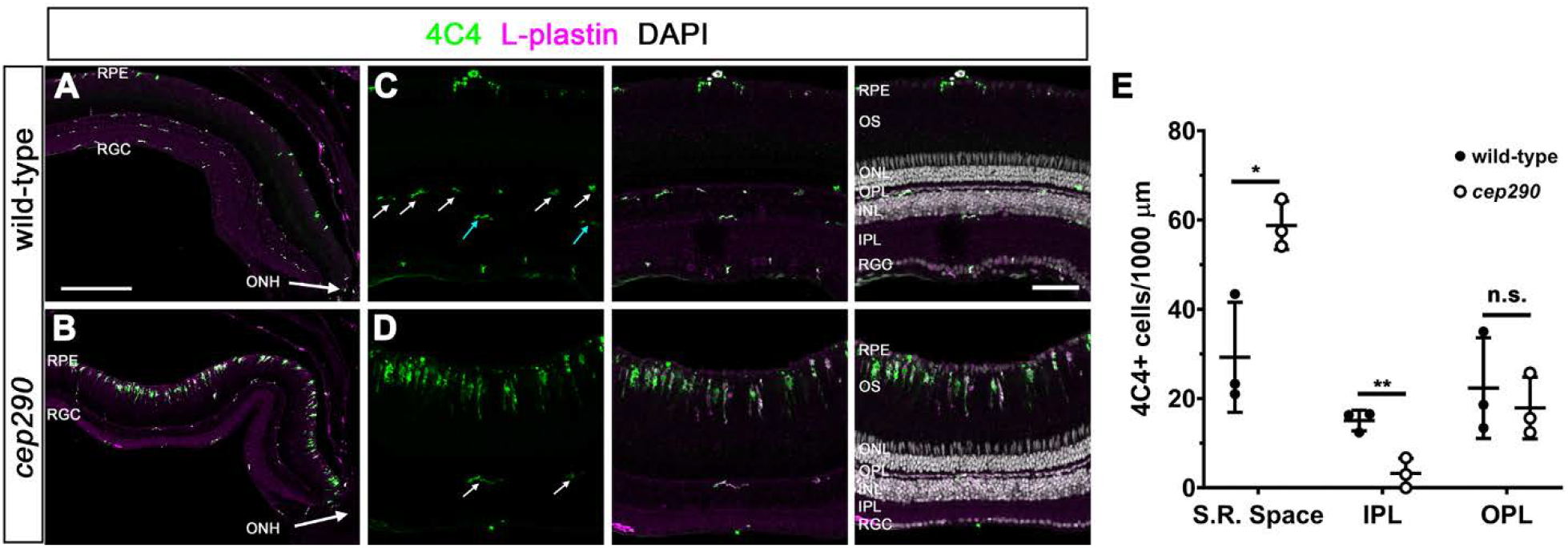
Immune cells accumulate in the subretinal space of *cep290* mutants. (A, B) Immunohistochemistry with the monoclonal antibody 4C4 (green) and anti-L-plastin (magenta) label microglia/macrophage in the dorsal retina of 6 mpf wild-type and *cep290* mutants. The optic nerve head (ONH) is located at the bottom right corner of each image. (C, D) Higher magnification images show the accumulation of activated microglia/macrophage in the subretinal space of *cep290* mutants. Ramified microglia/macrophage (4C4+/L-plastin+) were seen in the OPL (white arrows) of both wild-type and *cep290* mutants but only in the inner plexiform layer of wild-type retinas (cyan arrows). (E) Quantification of 4C4+ cells in different regions of the retina in 6 mpf fish (n = 3 per genotype). Data are plotted as means ± SD and p-values were generated by Welch’s t tests. \**p<0.05;* \*\**p<0.01*. RPE = retinal pigment epithelium, RGC = retinal ganglion cell layer, ONH = optic nerve head, OS = outer segments, ONL = outer nuclear layer, INL = inner nuclear layer, S.R. space = subretinal space, IPL = inner plexiform layer, OPL = outer plexiform layer. Scale bars: (A, B) 200 μm; (C, D) 50 μm.

### Transcriptome analysis of *cep290* mutant retinas

To gain a better understanding of transcriptional changes that occur during photoreceptor degeneration and that may promote regeneration, we performed RNA-seq analysis on retinas from 6 mpf *cep290* mutants (n = 4) and wild-type siblings (n = 4). Pairwise comparisons of all *cep290* mutant and wild-type transcriptomes revealed 1379 differentially expressed genes (DEGs; p_adj_ < 0.05; SI Appendix, Table S1). Of these, 235 DEGs were upregulated at least 2-fold (p_adj_ < 0.05). Pairwise comparisons of all samples were tested for enriched gene sets by Gene Set Enrichment Analysis (GSEA) and 11 pathways were upregulated in *cep290* mutants (Table 1; p ≤ 0.01; NES > 1.5; FDR < 0.1). The unfolded protein response, TP53, and apoptosis pathways were upregulated in *cep290* mutants, which likely reflects cellular stress and degeneration of photoreceptors. Upregulation of the interferon alpha, interferon gamma, IL-6/JAK/STAT3 signaling and TNFα signaling pathways were evidence of immune system activation in *cep290* mutants. These signaling pathways promote MG reprogramming and cell division following acute injury in healthy adult zebrafish (13, 63, 64), yet MG did not proliferate in *cep290* mutants. In injury models, Müller cell reprogramming involves TNFα-dependent expression and activation of the transcription factor Stat3 (13, 14, 65). Expression of *stat3* increases following retinal injury and suppression of *stat3* expression significantly inhibits regeneration (14). Activated phospho-Stat3 (p-Stat3) enters the nucleus and drives expression of reprogramming genes, including *ascl1a* and *lin28* (63, 64). In our RNA-seq dataset, expression of *stat3* was increased 1.86-fold (p < 0.0001) in *cep290* mutant retinas. To validate this result, we performed qRT-PCR on several factors essential for Müller cell reprogramming, including *tnfa, stat3, hmga1a*, and *yap1* (13, 15). As a positive control, we used the *Tg(rho:YFP-Eco.NfsB)gmc500* transgenic line that expresses a YFP-nitroreductase fusion protein in rod photoreceptors. Exposing this line to metronidazole (MTZ) results in a selective ablation of rods and robust MG activation (32, 66). MTZ-induced rod ablation resulted in a significant increase in the expression of all regeneration-associated genes (Figs. 2A-D). In *cep290* mutants, we confirmed a 2.1-fold increase (p < 0.05) of *stat3* and a 1.6-fold increase (p < 0.01) in *yap1* but detected no difference in expression of *tnfa* or *hmga1a* when compared to wild-type adults (Figs. 2A-D). The transcriptomic analysis suggests that photoreceptor degeneration stimulates inflammatory and upstream Stat3 signaling pathways, but this is insufficient to trigger MG reprogramming and proliferation.

**Figure 2.**
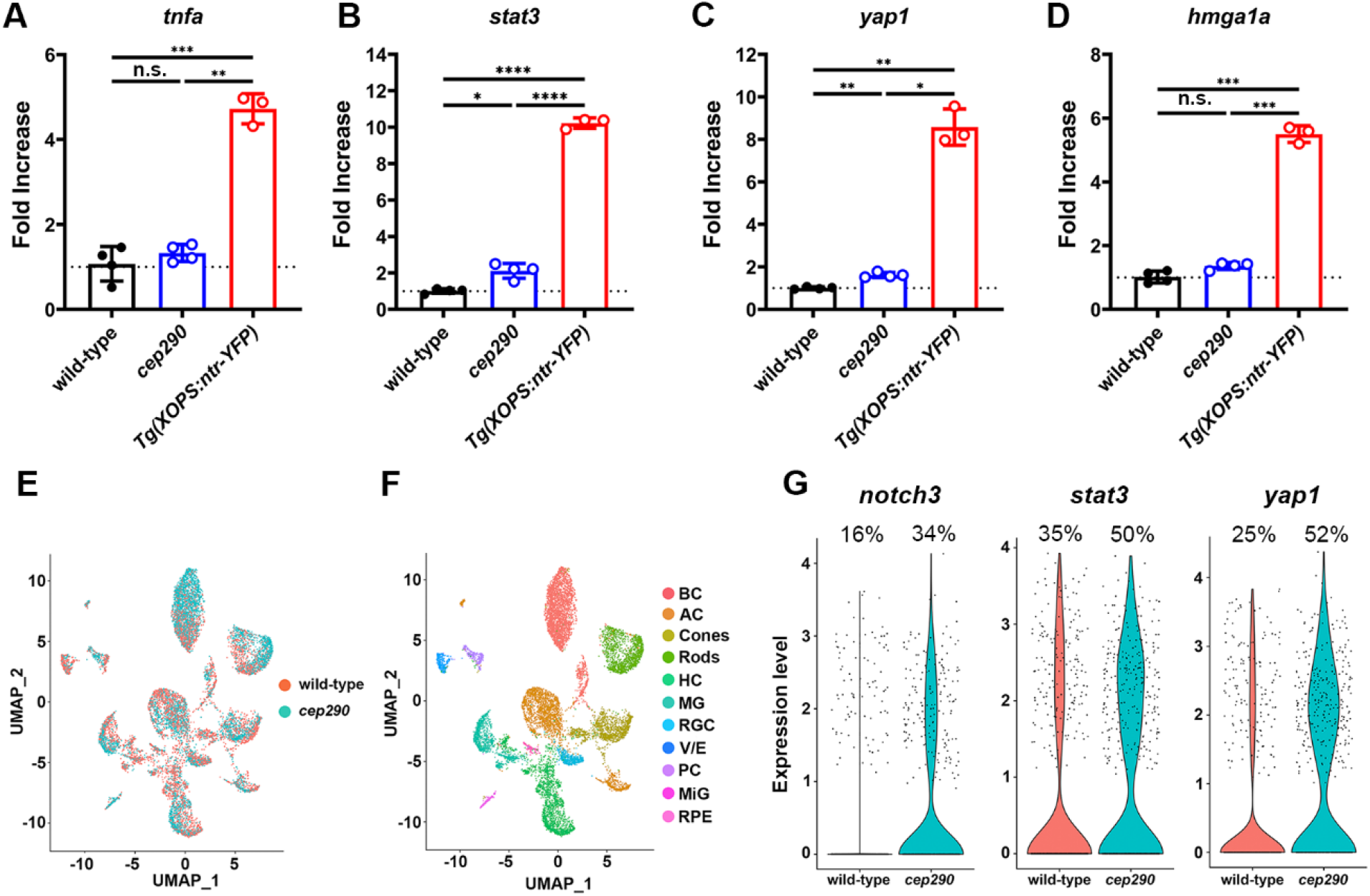
Expression of pro-regeneration genes was only modestly upregulated in 6 mpf *cep290* mutants. (A-D) qPCR for *tnfa, stat3, yap1*, and *hmga1a* in 6 mpf wild-type and *cep290* mutant zebrafish. Gene expression in MTZ-treated *Tg(rho:ntr-YFP)gmc500* zebrafish was used as a positive control for MG reactivity. Fold changes were calculated by the ΔΔC(t) method, with 18S rRNA used for normalization. No significant difference was observed between wild-type and *cep290* mutants for *tnfa* (*p* > 0.63) or *hmga1a* (*p* > 0.06). *cep290* mutants upregulated expression of *stat3* 2-fold (\**p* < 0.02) and *yap1* 1.6-fold (\*\**p* < 0.002) compared to wild-type animals. Expression of all genes was significantly upregulated in MTZ-treated *Tg(rho:ntr-YFP)gmc500* fish compared to both wild-type and *cep290* fish. Welch’s ANOVA with Dunnett T3 multiple comparisons test. \**p* < 0.02; \*\**p* < 0.002; \*\*\**p* < 0.0002; \*\*\*\**p* < 0.0001. (E) UMAP plot showing all cells obtained from sequencing with cells colored by sample. (F) UMAP plot showing identified cell types from sequencing data, including bipolar cells (BP), amacrine cells (AC), horizontal cells (HC), retinal ganglion cells (RGC), vascular/endothelial cells (V/E), pericytes (PC), microglia (MiG), and retinal pigment epithelium (RPE). (G) Violin plots showing expression of *notch3, stat3*, and *yap1* in MG between sample groups.

**Table 1.** GSEA Upregulated Pathways.

| **Gene Set** | **NES** | **FDR** |
| --- | --- | --- |
| Interferon alpha response | 2.38 | 0.000 |
| Interferon gamma response | 2.23 | 0.000 |
| IL-6/JAK/STAT3 signaling | 2.12 | 0.000 |
| Unfolded Protein Response | 1.76 | 0.007 |
| TP53 pathway | 1.70 | 0.012 |
| Myc targets | 1.59 | 0.030 |
| TNFα signaling via NFκB | 1.59 | 0.027 |
| Complement | 1.59 | 0.025 |
| Allograft rejection | 1.58 | 0.023 |
| mTORC1 signaling | 1.57 | 0.026 |
| Apoptosis | 1.54 | 0.029 |

NES = normalized enrichment score
FDR = false discovery rate

To determine if the observed transcriptomic changes are specific to MG, we performed scRNA-seq with *cep290* mutants and their heterozygote siblings (Figs. 2E, F). Differential expression analysis comparing MG from the two sample groups identified 561 significantly upregulated genes in *cep290* MG (SI Appendix, Table S2). Similar to whole retina RNA-seq, significantly more MG were observed to increase expression of *stat3* and *yap1* (Fig. 2G). Increased expression of *notch3* was also observed specifically in MG, which was not detected by whole retina RNA-seq. To build upon the scRNA-seq analysis, we compared the 561 DEGs to previously published transcriptomic data (15), which used pseudotime analysis of the regenerating retina to identify 3 MG states: resting, reactive, or proliferating. By comparing the 561 DEGs to known markers of each of these states, we identified 106 DEGs in *cep290* MG that were associated with any of these states. Of those 106 DEGs, 76 were associated with resting MG (SI Appendix, Fig. S2). This suggested an enrichment in rest-associated genes in *cep290* MG. We also compared the 561 DEGs to a predicted regulatory network within MG of the regenerating zebrafish retina (15). Of the 26 genes that belong to the predicted regulatory network (SI Appendix, Table S4), 21 belong to rest-associated modules (M2, M3, M4, M8; SI Appendix, Fig. S3) that were previously described (15).

To further assess Stat3 activation, the transgenic reporter line *Tg(gfap:stat3-GFP)mi35Tg* (63) was crossed into the *cep290* background. The *Tg(gfap:stat3-GFP)* line constitutively expresses *stat3-gfp* mRNA in MG but the unphosphorylated Stat3-GFP protein is degraded and remains undetectable in undamaged retinas (63). Following retinal injury, however, activated p-Stat3-GFP accumulates specifically in proliferating MG-derived progenitors. Animals were intraperitoneally injected with EdU to label proliferating cells and allowed to recover for 24 hrs. We assessed Stat3-GFP expression and proliferation using anti-GFP antibodies and EdU, respectively. In 6 mpf wild-type and *cep290* mutant retinas, Stat3-GFP was not detected and EdU^+^ cells were only observed in the ONL of *cep290* mutants (SI Appendix, Figs. S4A, B). As a positive control, animals were first injected with EdU and mechanically injured by a needle poke. Stat3-GFP accumulated at the site of injury and localized to EdU^+^ cells in both wild-type and *cep290* mutants (SI Appendix, Figs. S4C, D). We conclude that *cep290* mutants could activate Stat3 in response to acute injury. Surprisingly, the ongoing photoreceptor degeneration and inflammation observed in *cep290* mutants remains insufficient to activate a p-Stat3 that can be detected *in vivo*.

### Light-induced photoreceptor death stimulates regeneration in *cep290* mutants

To determine if regeneration by MG was possible in *cep290* mutants, we used high-intensity light to ablate photoreceptors (5, 67). Following light-induced photoreceptor cell death, MG dedifferentiate, re-enter the cell cycle, and produce neuronal progenitors that migrate to the ONL and differentiate to replace lost photoreceptors (5, 21, 67). To track proliferation of MG, the transgenic reporter line *Tg(gfap:eGFP)mi2002* (58) was crossed into the *cep290* background. At 48 hours post injury (48 hpi), animals were intraperitoneally injected with EdU and sacrificed at 72 hpi. In both wild-type and *cep290* mutant retinas at 72 hpi, EdU^+^ cells were observed in the INL and ONL (Figs. 3A, B). Within the INL, the EdU signal colocalized with GFP fluorescence from the *Tg(gfap:eGFP)* transgene, establishing the proliferating cells as MG. Importantly, there was no difference in the number of EdU^+^ cells in either the INL or ONL in light-damaged retinas from wild-type or *cep290* mutants (Fig. 3C). We next asked whether MG-derived progenitors in *cep290* mutants could regenerate photoreceptors following light damage. Photoreceptor degeneration and regeneration were assessed using the cone marker Zpr1 and peanut agglutinin (PNA), or the rod outer segment marker, Zpr3. Cell proliferation was measured by EdU incorporation. At 3 dpi, cones were ablated in light damaged wild-type and *cep290* mutants, and proliferation of neural precursors was observed in the INL (Figs. 4A, B, D, E). By 30 dpi, cones had regenerated in both wild-type and *cep290* mutants (Figs. 4C, F). Similarly, rod photoreceptors degenerated by 3 dpi but had regenerated by 30 dpi in both wild-type and *cep290* mutants (Figs. 4G-L).

**Figure 3.**
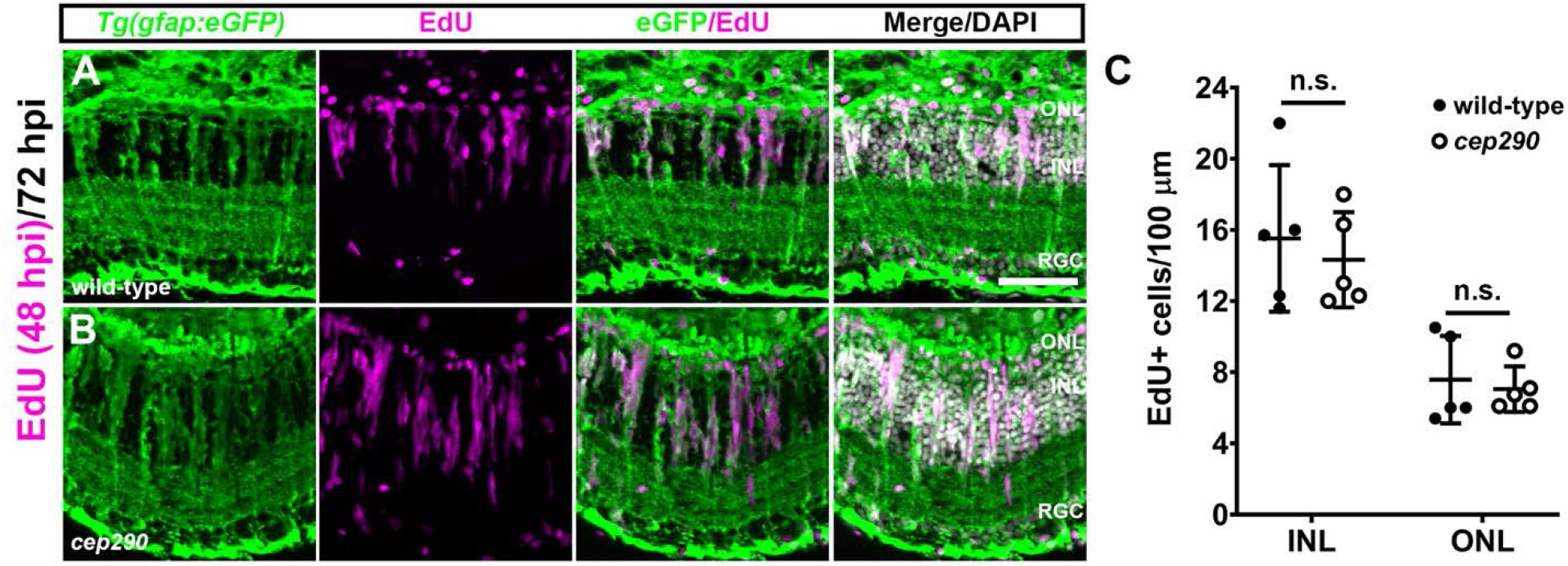
MG in *cep290* mutants respond to acute light damage. Retinal cryosections of 6 mpf wild-type (A) or *cep290* mutants (B) carrying the *Tg(gfap:eGFP)^mi2002^* transgenic reporter line were immunolabeled with anti-GFP (green) antibodies to visualize MG and processed for EdU labeling (magenta) to identify MG-derived progenitor cells at 72 hours post injury (hpi). (C) The number of EdU^+^ cells was not statistically different (n.s.) between wild-type and *cep290* mutants (INL: *p* > 0.99; ONL: *p* > 0.66; Mann-Whitney tests). Data are plotted as means ± SD. ONL = outer nuclear layer; INL = inner nuclear layer, RGC = retinal ganglion cell layer. Scale bar, 50 μm

**Figure 4.**
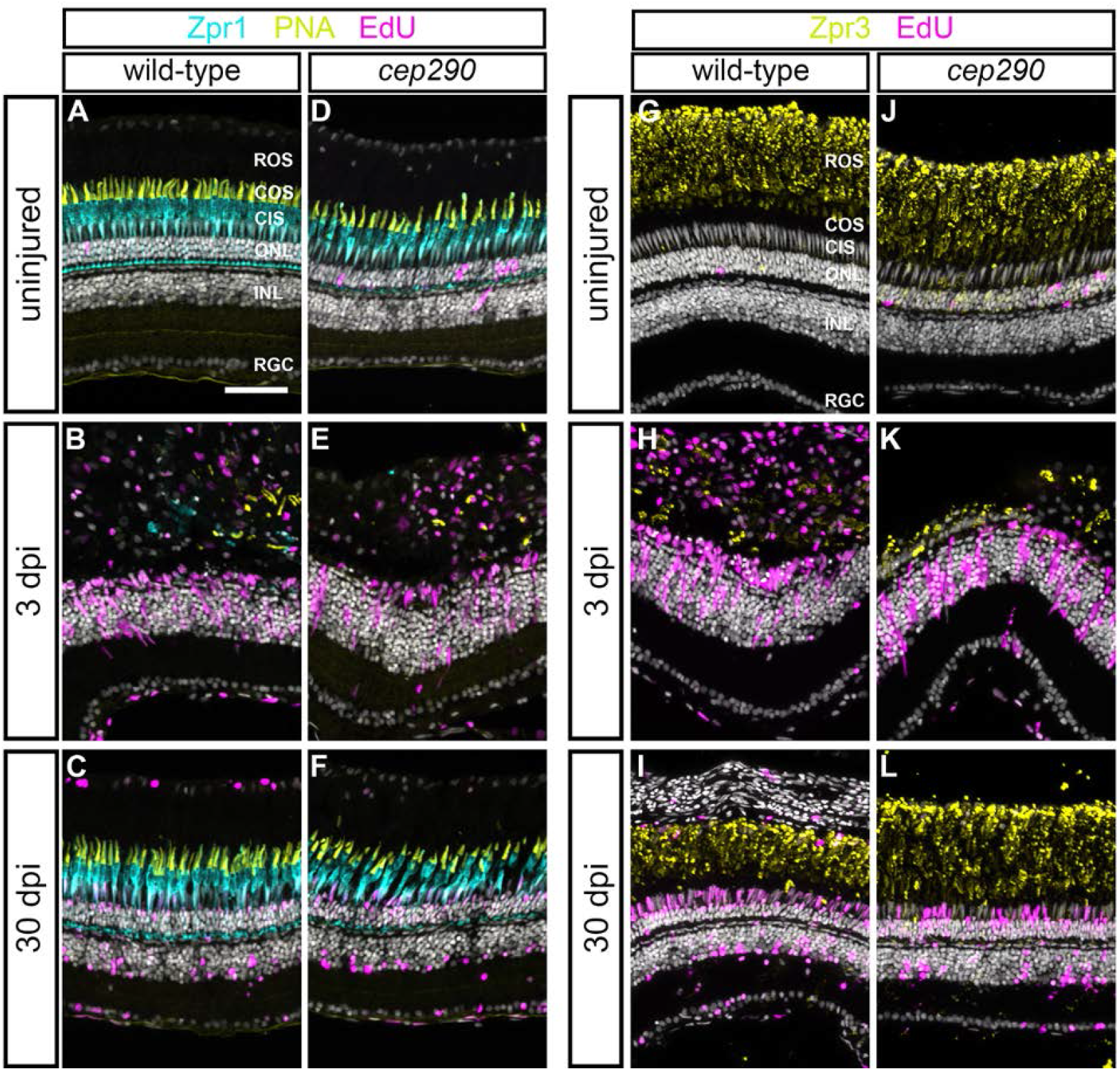
Photoreceptors regenerate in 6 mpf *cep290* mutants following acute light damage. (A-F) Views of cryosections of the dorsal retina of 6 mpf wild-type (A-C) or *cep290* mutants (D-F) immunolabeled with the red/green cone photoreceptor marker, Zpr1 (cyan), PNA (yellow) to visualize cone outer segments, and EdU (magenta) to detect cells that had undergone proliferation in undamaged retinas or at 3 or 30 dpi (days post injury). (G-L) Immunolabeling of retinas from 6 mpf wild-type (G-I) or *cep290* mutants (J-L) with the rod marker Zpr3 (yellow) and EdU (magenta) in undamaged retinas or at 3 or 30 dpi. ROS = rod outer segments; COS = cone outer segments; CIS = cone inner segments; ONL = outer nuclear layer; INL = inner nuclear layer, RGC = retinal ganglion cell layer. Scale bar, 50 μm.

Following light damage, regeneration can restore the cone density of wild-type retinas to pre-damage levels (25). Acute light damage triggers photoreceptor death in the central region of the dorsal retina, with the dorsal periphery and ventral retina areas largely spared from cell death and serving as an internal measure of pre-lesion cone densities (25). EdU incorporation during proliferation of neural progenitors marked the location of light damage in wild-type and *cep290* mutants, while few EdU^+^ cells were observed in the undamaged periphery (Figs. 5A, B). To determine if the density of regenerated cones in *cep290* mutant retinas was similar to that found in wild-type densities, we quantified cones in the damaged (“lesion”) and undamaged areas (“surround”) after 1 month of recovery in 3 mpf, 6 mpf, and 13 mpf fish. Cone density in the undamaged *cep290* retinas was consistently less than wild-type at 6 mpf and 13 mpf (Figs. 5C-E, blue bars). In both wild-type and *cep290* mutants regeneration restored cone density only to pre-damage levels (Figs. 5C-E, compare blue and red bars). That is, the total number of regenerated cones in *cep290* mutants remained lower than age-matched wild-type fish. These results indicate that zebrafish restore cones only to pre-lesion densities.

**Figure 5.**
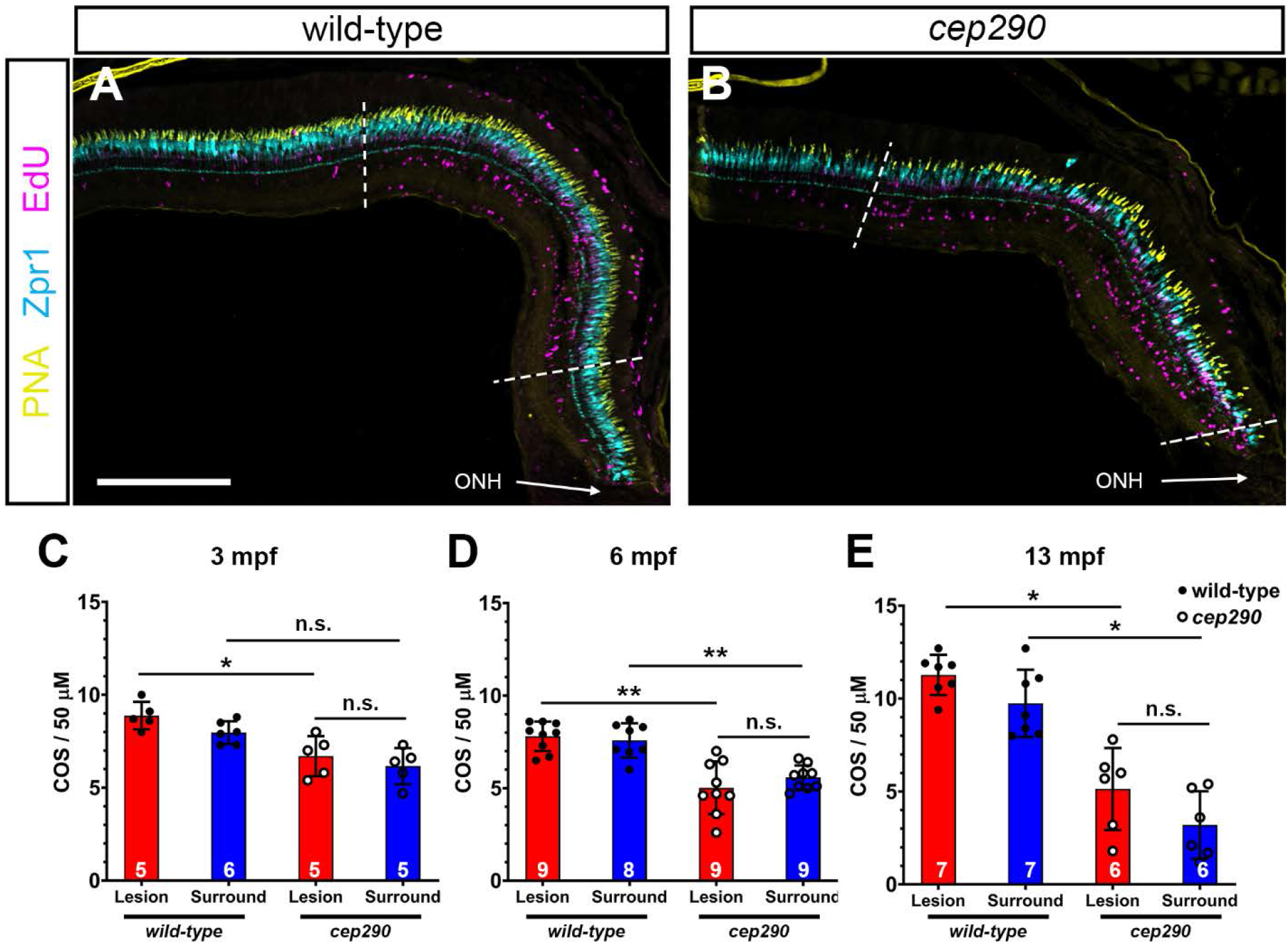
Cone regeneration is limited by pre-lesion cone density in *cep290* mutants. (A, B) Views of the dorsal retina in cryosections of 6 mpf wild-type and *cep290* mutant retinas at 30 dpi following light damage. Cryosections were immunolabeled with the red/green cone photoreceptor marker, Zpr1 (cyan), PNA (yellow) to visualize cone outer segments, and EdU (magenta) to detect cells that had undergone proliferation. Dotted lines indicate the regenerated area (i.e. lesion) based on highest density of EdU^+^ cells. (C-E) Quantification of cone outer segment density in the lesioned (red bars) and surrounding areas (blue bars) of wild-type (filled circles) and *cep290* retinas (open circles) at 30 dpi in animals aged 3 mpf, 6 mpf, and 13 mpf. Each data point represents quantification from an individual animal. Data are provided as means ± s.d. (3 mpf: \**p* < 0.05, Kruskal-Wallis test with Dunn’s multiple comparisons test; 6 mpf: \*\**p* < 0.005, Welch ANOVA test with Dunnett’s T3 multiple comparisons test; 13 mpf: \**p* < 0.05, Kruskal-Wallis test with Dunn’s multiple comparisons test). ONH = optic nerve head. Scale bar, 200 μm.

### Pharmacological modulation of glucocorticoid signaling impairs regeneration

Inflammation and microglia/macrophages are considered essential during the early proliferation phase of retinal regeneration in zebrafish (32, 35, 68). Recent reports found that pharmacological ablation of microglia/macrophages prior to injury inhibited proliferation of MG-derived progenitors (32, 36). Dexamethasone, a synthetic corticosteroid and agonist of the glucocorticoid receptor, is a potent anti-inflammatory agent. Pre-treatment with dexamethasone also inhibits MG proliferation following injury (32, 35). Conversely, blocking endogenous glucocorticoid receptor activity with the antagonist RU486 after NMDA-induced retinal degeneration increased proliferation MG-derived progenitor cells in the chick retina (69). Given these observations, we tested what effect treating *cep290* mutants with RU486 or dexamethasone had on microglia/macrophage reactivity, and the proliferation of Müller-glia derived progenitors or ONL rod precursors, respectively. We exposed 6 mpf animals to dexamethasone or RU486 and assessed microglia/macrophage accumulation and cell proliferation. As noted previously, the *cep290* mutants exhibit significantly more 4C4+ microglia/macrophages in the subretinal space compared to wild-type siblings (Figs. 6A, D; quantified in 6G). In both wild-type and *cep290* mutants, exposure to dexamethasone significantly reduced the number of 4C4+ microglia/macrophages in the subretinal space and in the inner retina as compared to control animals exposed to vehicle (Figs. 6A-H). Exposure to RU486 had no effect on the number of microglia/macrophage in either the subretinal space or the inner retina (Figs. 6G and H), consistent with the findings that RU486 had no effect on microglia/macrophage reactivity in chick (69). Dexamethasone treatment also significantly reduced the number of proliferating cells in the ONL of *cep290* mutants (Figs. 6L-O). Although dexamethasone appeared to reduce the number of PCNA+ cells in the ONL of wild-type retinas, the results were not statistically significant, as few PCNA+ cells are found in undamaged wild-type retinas. As expected, treatment with RU486 had no effect on the number of PCNA+ cells in wild-type retinas. However, RU486 also had no effect on the number of PCNA+ cells found in the ONL or INL of *cep290* mutants (Figs. 6O and 6P). This is in contrast to previous findings in chick where RU486 treatment increased the number of proliferating MG-derived progenitors nearly six-fold (69). These results suggest that immunosuppression inhibits proliferation of rod precursors in *cep290* mutants but that inhibition of the glucocorticoid receptor– i.e., blocking endogenous anti-inflammatory activity – is insufficient to stimulate proliferation of either MG-derived retinal progenitors or rod precursors in *cep290* mutants.

**Figure 6.**
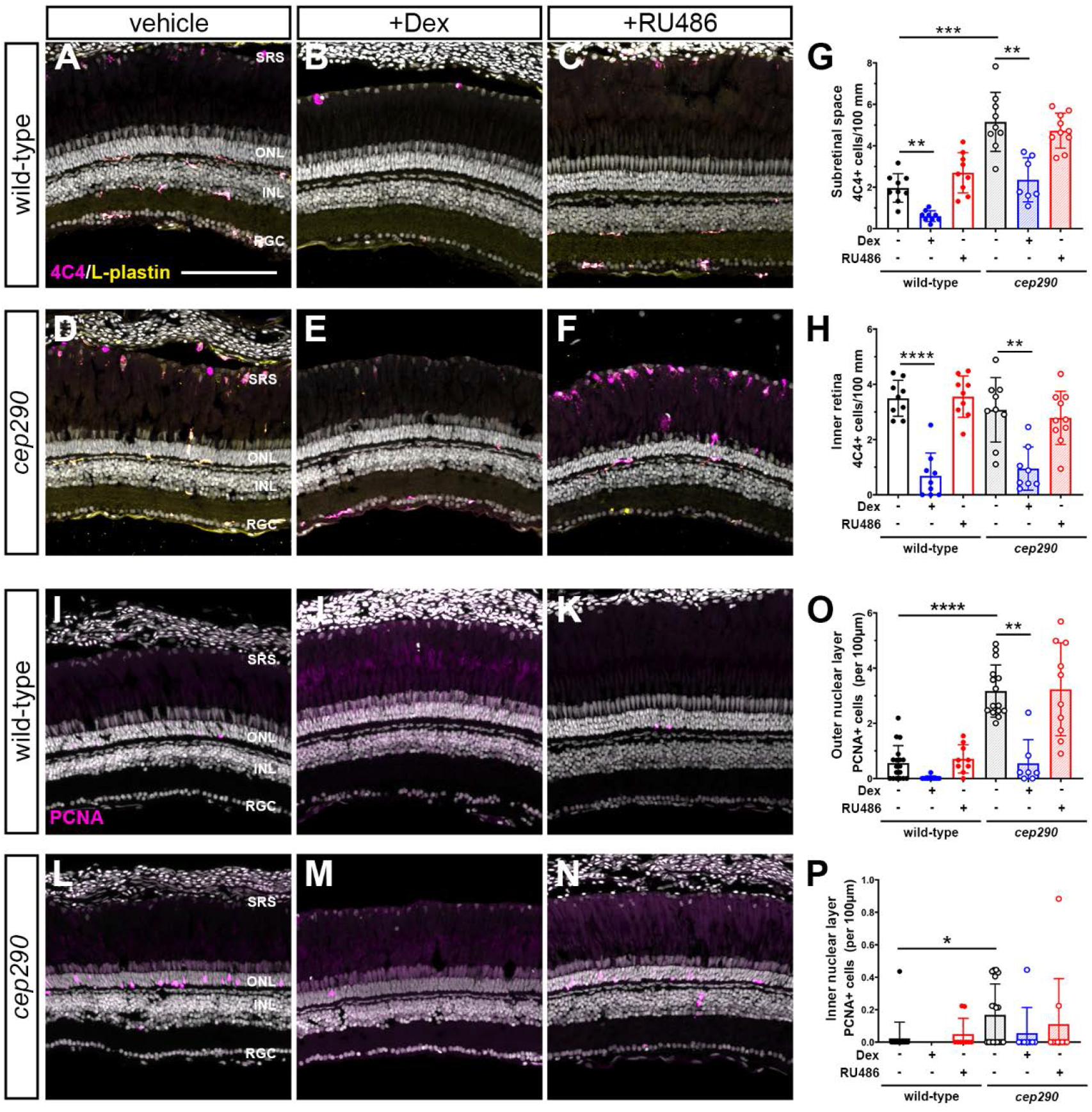
Immunosuppression with dexamethasone inhibits proliferation and microglia/macrophage activation in *cep290* mutants. (A-F) Immunohistochemistry of the dorsal retina with markers for microglia/macrophage (4C4, magenta) and leukocytes (L-plastin, yellow) in 6 mpf wild-type and *cep290* mutant animals treated with methanol (vehicle/left column), dexamethasone (middle column) or RU486 (right column). (G, H) Quantification of 4C4+ cells in the subretinal space and inner retina, respectively of wild-type (filled circles/open bars) and *cep290* mutants (open circles/hashed bars). (I-N) Immunohistochemistry of the dorsal retina with PCNA (magenta) in 6 mpf wild-type and *cep290* mutant animals. (O, P) Quantification of PCNA+ cells in the outer nuclear layer and inner nuclear layer, respectively, of wild-type (filled circles/open bars) and *cep290* mutants (open circles/hashed bars). Counts following dexamethasone treatment are plotted in blue, whereas counts following RU486 treatment are plotted in red. Each data point represents counts from the dorsal retina of one eye. For graphs in G and H, the significance in differences was determined using Welch ANOVA test with Dunnett’s T3 multiple comparisons test (\*\**p* < 0.01, \*\*\**p* < 0.001, \*\*\*\**p* < 0.0001). For graphs in O and P, the significance in differences was determined by Kruskal-Wallis test with Dunn’s multiple comparisons test (\**p* < 0.05, \*\**p* < 0.01, \*\*\*\**p* < 0.0001). SRS = subretinal space; ONL = outer nuclear layer; INL = inner nuclear layer, RGC = retinal ganglion cell layer. Scale bar, 100 μm.

### Chronic immunosuppression rescues cone photoreceptors

We next asked whether sustained immunosuppression could improve cone survival. While the immune response appears critical for regeneration, chronic inflammation also contributes to ongoing neuronal degeneration. When dexamethasone was provided after injury, cone survival was enhanced in *mmp9^-/-^* zebrafish mutants (35). Furthermore, genetic ablation of microglia in the *rd10* mouse improved long-term survival of photoreceptors (28). We hypothesized that although sustained immunosuppression may inhibit rod cell regeneration, it may prolong cone photoreceptor survival in *cep290* mutants. To test this, we utilized the zebrafish *irf8* mutant line (70). The transcription factor interferon regulatory factor 8 (IRF8) is essential for the development of all embryonic macrophages and microglia in zebrafish (70). Adult *irf8* mutants are viable and lack most resident microglia in the CNS (71). Quantification of microglia/macrophage in flat-mounted retinas found a 92.8% reduction in microglia/macrophage density in *irf8* mutants compared to wild-type siblings (Fig. S5A-C). We next generated *cep290;irf8* double mutants and compared the phenotypes to *cep290* and *irf8* single mutants at 6 mpf. Whereas healthy microglia/macrophages were observed in the plexiform layers of wild-type animals (Fig. 7A) and reactive microglia/macrophages accumulated in the subretinal space of *cep290* mutants (Fig. 7B), there was a striking absence of microglia/macrophages the retinas of either *irf8* mutants or the *cep290;irf8^-^* mutants (Figs. 7C, D). To assess proliferation, 6 mpf animals were injected with EdU and collected two days post injection (2 dpi) and the retinas were examined by immunohistochemistry. While proliferation of rod precursors in the ONL was minimal in wild-type animals (Fig. 7E) and significantly greater in *cep290* mutants (Fig. 7F), few proliferating cells were observed in the *irf8* mutants (Fig. 7G). Consistent with data showing that immunosuppression impaired regeneration, the number of proliferating ONL cells in *cep290;irf8* mutants significantly lower than in *cep290* mutants but not statistically different from wild-type or *irf8* mutants (Figs. 7E-H and 7M). Finally, we quantified cone density in 6 mpf animals. Compared to wild-type siblings, the number of cones was reduced in *cep290* mutants but not *irf8* mutants, which indicated that loss of *irf8* does not impair cone survival (Figs. 7I-K, 7N). When cone density was quantified in *cep290;irf8* mutants, the number of cones was significantly greater than *cep290* mutants but not statistically different than wild-type siblings (Figs. 7L, N). These results indicate that immunosuppression can prolong cone survival in *cep290* mutants. The lack of proliferation in the INL or ONL of the *cep290;irf8* mutants also indicates that the increased number of cones is most likely due to cone preservation and not enhanced regeneration. Taken together, these data indicate that inflammation stimulates ONL cell proliferation in *cep290* mutants to maintain rod cell numbers, but that the absence of microglia/macrophage promotes cone survival in *cep290* mutants.

**Figure 7.**
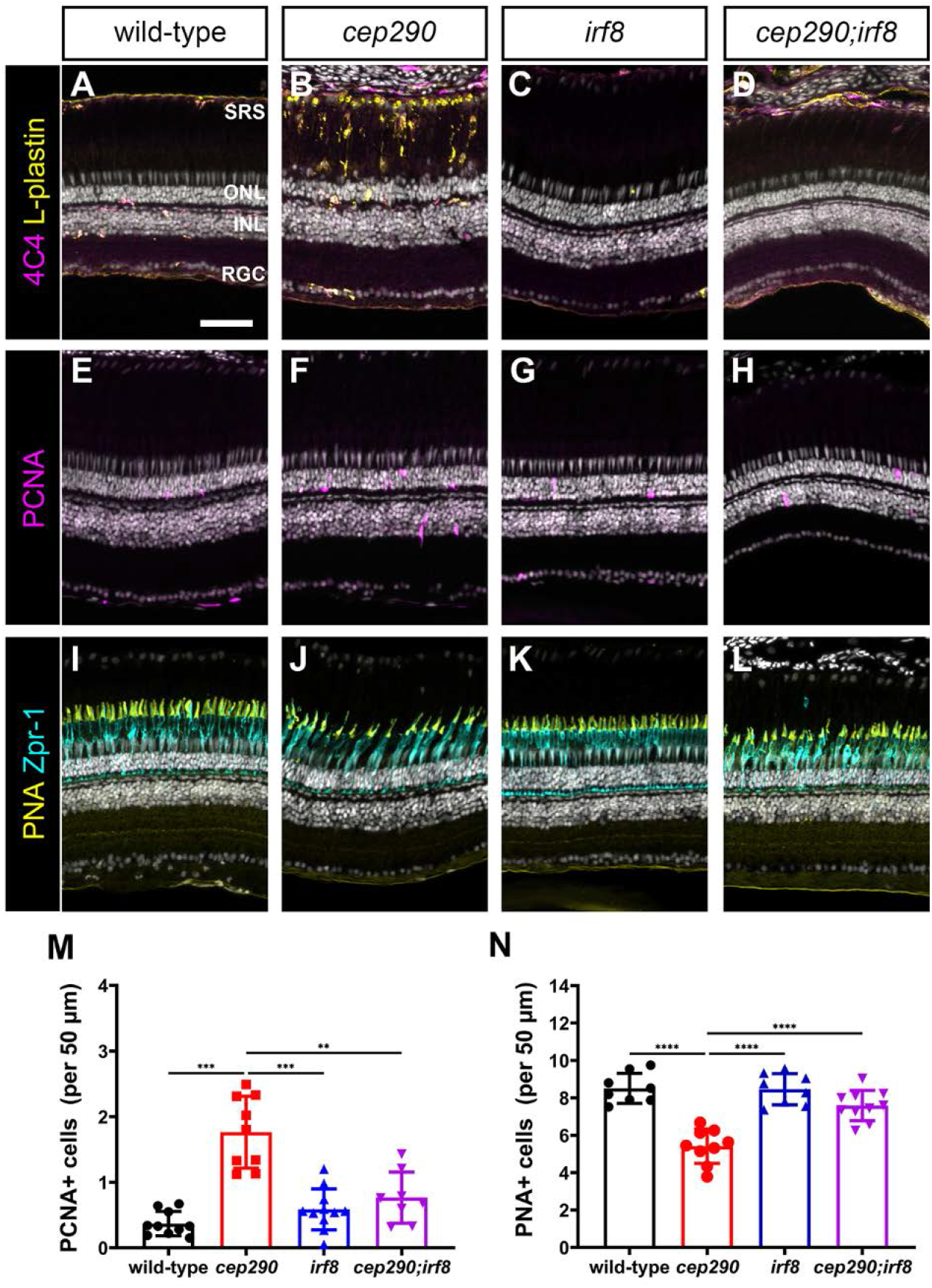
Mutation of *irf8* reduces the number of activated microglia/macrophage and proliferating cells and promotes cone survival in *cep290* mutants. (A-D) Immunohistochemistry of the dorsal retina with markers for microglia/macrophage (4C4, magenta) and leukocytes (L-plastin, yellow) in 6 mpf wild-type, *cep290* mutants, *irf8* mutants, and *cep290;irf8* mutants. (E-H) Immunohistochemistry of the dorsal retina at 3 days post-injection with EdU (magenta) to mark proliferating cells in 6 mpf wild-type, *cep290* mutants, *irf8* mutants, and *cep290;irf8* mutants. (I-L) Immunohistochemistry of the dorsal retina with markers for cone inner segments (Zpr-1, cyan) and cone outer segments (PNA, yellow) in 6 mpf wild-type, *cep290* mutants, *irf8* mutants, and *cep290;irf8* mutants. (M, N) Quantification of PCNA+ cells in the outer nuclear layer and PNA+ outer segments in wild-type (filled black circles) and *cep290* mutants (filled red circle), *irf8* mutants (filled blue triangles), and *cep290;irf8* mutants (filled purple triangles). Each data point represents counts from the dorsal retina of one eye. For the graph in M, the significance in differences was determined using Welch ANOVA test with Dunnett’s T3 multiple comparisons test (\*\**p* < 0.01, \*\*\**p* < 0.001). For the graph in N, the significance in differences was determined by one-way ANOVA with Tukey’s multiple comparisons test (\*\*\**p* < 0.001, \*\*\*\**p* < 0.0001). SRS = subretinal space; ONL = outer nuclear layer; INL = inner nuclear layer, RGC = retinal ganglion cell layer. Scale bars, (A, B) 400 μm; (D-O) 50 μm.

### Notch inhibition induces MG to re-enter the cell cycle in *cep290* mutants

In zebrafish, Notch signaling suppresses regeneration by maintaining MG in a quiescent state (72–74) and proliferation requires the downregulation of *notch3* expression in MG (75, 76). As more MG in *cep290* mutants maintain expression *notch3* (Fig. 2G), we hypothesized that persistent Notch signaling maintained MG quiescence in *cep290* mutants despite chronic degeneration. To inhibit Notch signaling, adult *cep290* mutants and wild-type siblings were intraperitoneally injected with the γ-secretase inhibitor RO4929097 or 1% DMSO (vehicle) every 12 hours for 4 days and collected 24 hr after the last injection. Proliferation was assessed by PCNA staining (Fig. 8). In DMSO-control retinas, very few PCNA+ cells were observed in wild-type animals (Fig. 8A), while PCNA+ cells were limited primarily to the ONL in *cep290* mutants (Fig. 8B). Following RO4929097 treatment, a significant increase in PCNA+ cells were observed in the INL of both wild-type and *cep290* mutant retinas (Figs. 8C, D). Importantly, Notch inhibition resulted in a significantly greater number of PCNA+ cells in the INL of *cep290* mutants than in wild-type retinas (Fig. 8E), roughly matching the 2-fold increase in the number of *notch3*-expressing MG observed in *cep290* mutants by scRNA-seq (Fig. 2G). The PCNA+ cells were often observed in clusters within the INL of *cep290* mutants rather than the single PCNA+ cells observed in wild-type retinas, consistent with proliferation of MG and closely associated MG-derived progenitor cells. This is similar to the synergistic stimulation of proliferation seen following co-injection of RO4929097 and TNFα in the uninjured retina (72). Interestingly, Notch suppression had no effect on the proliferation of rod precursors in the ONL (Fig. 8F). When combined with our transcriptomic data, these results suggest that the regenerative response of MG in *cep290* mutants is suppressed by Notch signaling but that repressing Notch signaling in the chronic degenerative/inflammatory environment of the *cep290* retina is sufficient to induce MG proliferation.

**Figure 8.**
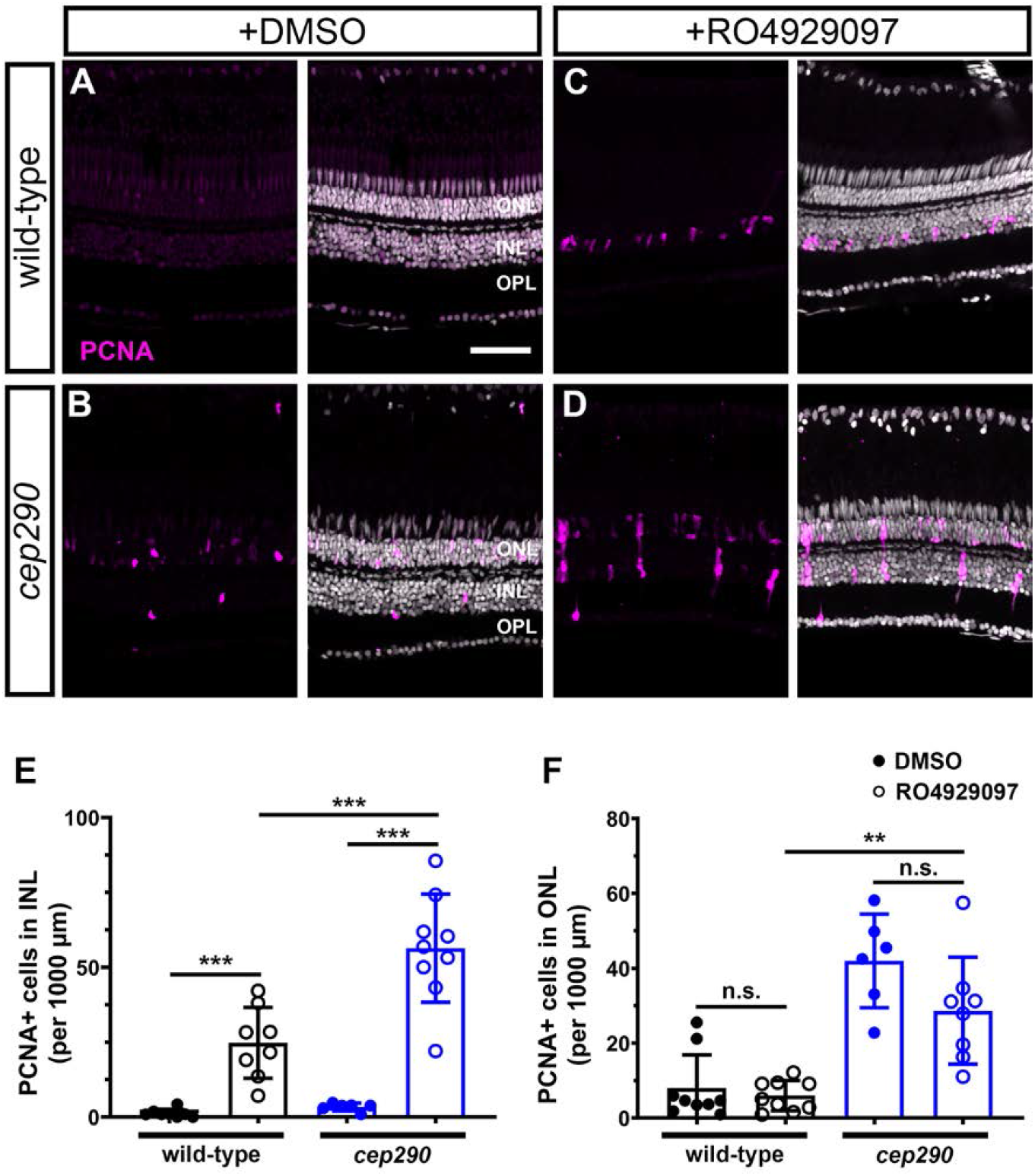
Inhibition of Notch signaling by injection of RO4929097 induces more robust Muller glia proliferation in 6 mpf *cep290* mutant retinas than in wild-type retinas. Wild-type (top row) and *cep290* mutant fish (bottom row) were injected intraperitoneally with 1% DMSO or 1 mM RO4929097 for 4 consecutive days. (A, B) Immunofluorescent images of dorsal retinas from DMSO injected fish or (C, D) RO4929097-injected fish stained with PCNA (magenta). (E) Quantification of PCNA+ cells in the INL or (F) ONL of DMSO-treated wild-type (filled black circles) and *cep290* mutants (filled blue circles) or RO4929097-treated wild-type (open black circles) and *cep290* mutants (open blue circles). Each data point represents counts from the dorsal retina of one eye and graphs represent mean ± s.d. Significance in differences were determined using Mann-Whitney tests except for comparisons between RO4929097-treated wild-type to *cep290* mutants, which used T-test with Welch’s correction (\*\**p* < 0.005; \*\*\**p* < 0.001). ONL = outer nuclear layer; INL = inner nuclear layer, OPL = outer plexiform layer. Scale bar: 50 μm.

## Discussion

Strategies that aim to stimulate endogenous repair in the retina are an attractive therapeutic approach for progressive retinal degenerative diseases. Identifying both the extrinsic cues and the intrinsic molecular mechanisms necessary for MG reprogramming and proliferation, and which regulate the differentiation of MG-derived retinal progenitors, remain central to achieving these goals. Much of the foundational knowledge on retinal regeneration has come from studying how zebrafish respond to acute damage (68, 77). Interestingly, several zebrafish mutants exhibiting progressive photoreceptor degeneration do not initiate a robust regenerative response to chronic cell loss (55, 57, 78). Here, characterization of the zebrafish *cep290* mutant, which undergoes progressive photoreceptor degeneration, has revealed that the molecular pathways required for Muller cell reprogramming remain intact in the diseased retina and helped refine the views of inflammation in driving regeneration. Specifically, we report that slow, progressive photoreceptor degeneration results in a sustained immune response exemplified by reactive microglia/macrophages and upregulation of inflammation-related genes in *cep290* mutants. Further, we show that immune suppression protects cone cells from degeneration, and that sustained expression of Notch signaling and quiescence-associated genes in MG prevents the re-entry into the cell cycle even in the presence of chronic inflammatory signals. This study is the first to show that in response to chronic degeneration, zebrafish MG adopt a novel state characterized by the simultaneous expression of genes associated with quiescence and reactivity.

Like the response to photoreceptor degeneration in mammals, *cep290* mutant retinas showed evidence of microglia/macrophage activation and increased inflammation, as well as an absence of a MG-based regenerative response. The immune cells accumulated in the subretinal space and the number of immune cells in the INL was reduced, similar to what has been observed in mouse models of RP (28, 79). By using RNA-seq to explore whole-retina gene expression, we noted the upregulation of several pathways associated with inflammation, including interferon alpha, interferon gamma, and Jak/Stat3 signaling. We also noted the upregulation of the recruitment factors for neutrophils (*cxcl8, cxcl18a. 1, cxcl18b*) and macrophages (*il34*), all of which were also upregulated in a zebrafish model of RPE damage (80). These data suggest that a similar inflammatory response exists to both chronic degeneration and acute damage within the zebrafish retina. Given the observed immune response, it is surprising that MG did not proliferate in *cep290* mutants. Several studies have documented a strong correlation between inflammation and MG proliferation following retinal cell death (32–36, 81). In fact, injection of the inflammatory cytokine TNFα into undamaged retinas is sufficient to induce proliferation (72). In zebrafish lacking *matrix metalloproteinase-9* (*mmp-9*), TNFα expression was elevated over wild-type in both undamaged and damaged retinas and acute light injury triggered an overproduction of MG-derived progenitors in *mmp-9* mutants (35). Dexamethasone, a potent anti-inflammatory agent, suppresses MG proliferation following injury (32, 35). Ablation of microglia by the drug PLX3397 also reduces proliferation of MG following laser injury (36). Our results indicate, however, that inflammation alone was insufficient to induce MG proliferation in a model of progressive photoreceptor degeneration. The loss of *cep290* function was likely not a factor as photoreceptor death also fails to induce MG proliferation in the *bbs2* mutant (55) or the *arylhydrocarbon interaction protein like 1b* (*aipl1b*) mutant (56). Furthermore, acute light damage triggered MG proliferation in both *cep290* and *bbs2* mutants, indicating that both mutants could respond to acute damage. The results from scRNA-seq analysis suggest that sustained expression of *notch3* in MG of *cep290* mutants inhibits proliferation even in a chronic inflammatory environment. MG maintain an “injury-response threshold” to limit regeneration and this threshold is controlled by Notch signaling (82). Several studies have confirmed that Notch signaling maintains MG quiescence (72–74, 76). Following acute injury, expression of *notch3* is rapidly downregulated to permit expression of regeneration-associated genes (76). Notch3 reportedly signals through the transcriptional repressor Hey1 to limit chromatin accessibility and establish the injury threshold. Morpholino knockdown of either *notch3* or *hey1* significantly increases MG proliferation following injury (76, 82). The link between inflammation and Notch inhibition remains unclear. TNFα expression and RO4929097-mediated Notch suppression act synergistically to induce MG proliferation but it is not known if these factors are mechanistically linked or functionally independently. Why MG in *cep290* mutants maintain Notch3 expression in spite of inflammatory signals remains unknown. Additional studies are also needed to identify the factors that inhibit Notch following acute injury.

Our results also show that proliferation of rod precursors depends on inflammation but not Notch signaling. The rod precursors are derived from MG and are maintained as a population of slowly dividing unipotent stem cells in the ONL that proliferate rapidly in response to rod death (22, 27). Our data show that the proliferating cells in the ONL of *cep290* mutants differentiate exclusively into rods, confirming them as rod precursors. We demonstrated that immunosuppression with dexamethasone or with an *irf8* mutant impaired rod precursor proliferation, indicating that rod precursors respond to inflammatory signals. Interestingly, injection of RO4929097 did not impact rod precursor proliferation. A previous report found that morpholino-induced knockdown of *notch3* did not increase the number of PCNA+ cells within the ONL at 96 hr following light damage (76). Taken together, these results suggest that Notch signaling does not influence the behavior of rod precursors.

Inflammatory signals enhance the regenerative response to retinal damage in zebrafish (13, 35, 56) but studies also show that inflammation potentiates photoreceptor degeneration in mouse models of retinitis pigmentosa (28, 38) and inflammatory signals from microglia limit regeneration in mammals (83). As immunomodulation has been posited as a potential therapy for retinitis pigmentosa (37, 38), we investigated the effects of chronic immunosuppression on photoreceptor regeneration and survival. When retinal microglia/macrophages were depleted in *cep290;irf8* double mutants, cone photoreceptors were protected from degeneration. Although rod precursor proliferation was reduced in *cep290;irf8* double mutants, we found no evidence of accelerated rod degeneration, suggesting that rods were also spared. This was unexpected as loss of Cep290 leads to cell-autonomous photoreceptor death by destabilizing the architecture and function of the connecting cilium (84). Lowering inflammation may mitigate the cellular stress associated with trafficking defects and slow photoreceptor death. Genetic and pharmacological inhibition of microglia function was reported to improve photoreceptor survival in mouse IRD models (28, 85–88). Cone survival following regeneration also requires resolution of the inflammatory response, suggesting that prolonged inflammation exacerbates photoreceptor degeneration (35). It is reasonable to speculate that prolonged inflammation may partially explain why cones only regenerate to pre-lesion densities following light damage in *cep290* and *bbs2* models. Perhaps the inability to resolve higher levels of inflammation limits cone survival following light damage. Alternatively, chronic inflammation could decrease survival of MG-derived progenitors.

The goal of studying zebrafish models of retinal degeneration is to determine how the mechanisms that promote regeneration function in different contexts. In zebrafish, both acute injury and progressive degeneration activate microglia/macrophages and increase inflammation (32–34, 55). In contrast, microglia activation appears to inhibit MG-associated regeneration following acute injury in mice (83). The role of Notch is also complex and context-dependent. The Notch pathway remains active in MG following damage in the mouse (15) and in zebrafish with progressive degeneration (this study), but Notch is downregulated in zebrafish MG following widespread acute injuries (15, 76). Furthermore, Notch inhibition combined with overexpression of the key reprogramming factors *Ascl1a* and *Lin28* did not exert the same pro-regenerative response in mouse MG as was observed following similar expression studies in zebrafish (89). Developing strategies for successful regeneration of photoreceptors in diseases like retinitis pigmentosa or age-related macular degeneration will require a better understanding of how inflammation and Notch signaling function in both injury and disease states.

## Materials and Methods

### Animal maintenance

Adult zebrafish were maintained and housed in 1.5 L, 3.0 L, and 10 L tanks in an Aquatic Habitats recirculating system (Pentair; Apopka, FL, USA) with a 14:10 light-dark cycle with ambient room lighting. Zebrafish lines utilized in this study included the mutant lines *cep290^fh297^* (1) and *irf8^st95^* (2), the transgenic reporter lines *Tg(Xla.Rho.eGFP)fl1* (3), *Tg(gfap:eGFP)mi2002* (4), and *Tg(gfap:stat3-eGFP)mi35* (5), as well as the transgenic line *Tg(rho:YFP-Eco.NfsB)gmc500* (herein referred to as *Tg(rho:ntr-YFP)*) to ablate rod photoreceptors (6). Animals from heterozygous crosses of *cep290* or *irf8* lines were genotyped using high resolution melt analysis (HRMA) or PCR using specified primer sets (Table S5). Homozygote, heterozygote, and wild-type siblings were identified for each cross. Transgenic lines were confirmed by PCR using primers specific for GFP (Table S5). Fin clips were added to 50 microliters of a 50 mM NaOH solution heated to 95 °C for 10 min and then neutralized with 5 microliters of 1M Tris pH 8.0 before cooling to room temperature. Experiments included animals of both sexes and were used at the specified ages. All animal procedures were done with approval by the Institutional Animal Care and Use Committee (IACUC) at the Cleveland Clinic and in accordance with relevant guidelines and regulations, including the ARVO statement for the use of animals in research.

### Light Damage

Light damage experiments were performed using a protocol adapted from Thomas and Thummel (7). Adult zebrafish were first dark adapted for 36-42 hours. Up to 5 animals were placed in a 250 mL glass beaker with system water that was seated inside a 1L glass beaker with Milli-Q water and exposed to high-intensity light from a 120W X-CITE series 120Q metal halide lamp (Excelitas) for 30 minutes and then exposed to 14,000 lux light from an illumination cage for 50 minutes. Animals were allowed to recover in system water for either 3 days or 30 days. To label proliferating cells during early stages of regeneration, animals were injected intraperitoneally with 20 μL of a 20 mM EdU solution (PBS) at 2 days post injury and eyes were enucleated 24 hours later (3 days post injury). To assess regeneration, animals were allowed to recover for 30 days post injury prior to enucleation.

### Immunohistochemistry

Adult zebrafish were deeply anesthetized with tricaine methanesulfonate (0.4 mg/mL) and decapitated with a razor blade. Eyes were rapidly enucleated, immersed for two hours in 4% paraformaldehyde and washed in phosphate buffered saline (PBS). Samples were equilibrated in 5% sucrose/PBS for 3 hrs at room temperature and transferred to 30% sucrose/PBS overnight at 4 °C. Eyes were washed in a 1:1 solution of 30% sucrose:Tissue Freezing Medium overnight at 4 °C and embedded for cryosectioning.

Transverse cryosections sections (10 μm) were cut and mounted on Superfrost Plus slides and dried at room temperature overnight. Slides were washed 3×10 min in PBS and then incubated in blocking solution (PBS + 2% BSA, 5% goat serum, 0.1% Tween-20, 0.1% DMSO) for 1 hr. The following primary antibodies were used: mouse monoclonal zpr1 (1:100, Zebrafish International Resource Center (ZIRC), Eugene, OR, USA), mouse monoclonal zpr3 (1:100, ZIRC), mouse monoclonal 4C4 (1:1000, a gift from Dr. Peter Hitchcock, University of Michigan), rabbit polyclonal L-plastin (1:1000, GeneTex, Irvine, CA, USA, GTX124420), mouse monoclonal PCNA (1:100, Sigma, St. Louis, MO, USA, clone PC-10), peanut agglutinin (PNA)-lectin conjugated to Alexa-568 (1:100, ThermoFisher, Waltham, MA, USA). EdU labeling was detected with the Click-iT Edu Alexa Fluor-555 Imaging Kit (ThermoFisher). Alexa-conjugated secondary antibodies were used at 1:500 in blocking buffer and incubated for at least 1 hr. Slides were counterstained with 4,5-diamidino-2-phenylendole (DAPI) to stain nuclei.

### Image acquisition and quantification

Z-stacked images of 5-15 μm immunostained cryosections were imaged on a Zeiss Imager Z.2 equipped with an ApoTome using 10x dry, 20x dry, or 63x oil immersion lenses (Zeiss). Images were acquired with Zen2 software and post-processed in ImageJ. All imaging and quantitative analysis was performed on dorsal retina sections, which contained or were immediately adjacent to the optic nerve. Cells or labels of interest (e.g. PNA, EdU, PCNA) were manually quantified in maximum-projection z-stacks of the dorsal retina and densities or ratios were calculated by measuring the curvilinear distance of retina in each section using ImageJ. Each data point represents the ratio or density from the dorsal region of a central retinal section from one eye.

### Lineage Tracing

Retinal sections from *Tg(XOPS:eGFP); cep290* fish that were injected with EdU 4 weeks prior to sacrifice were immunostained with zpr1 and imaged with a Zeiss Imager.Z2 with Apotome.2 attachment. Labeled cells were identified in single optical sections. Using DAPI staining as a guide, ROIs were drawn around the nucleus of cells labeled with either zpr1 or EdU. The GFP fluorescence in each ROI was normalized to the average GFP fluorescence in the nuclei of 5 random GFP+ cells in the ONL from the same image. Thus, rods will have a fluorescence intensity near 1, whereas cones will have an intensity near 0. As a negative control, we also measured background GFP fluorescence of random nuclei in the INL, which does not contain photoreceptors.

### Metronidazole treatment

To ablate photoreceptors using the *Tg(rho:ntr-YFP)* line, adult zebrafish were incubated in system water containing 10 mM metronidazole (Sigma) for 24 hrs in a 28 °C incubator. Animals were transferred to fresh system water and dark-adapted for 18 hours prior to euthanasia. The eyes were removed and retinas were dissected as described above.

### Dexamethasone and RU486 treatments

Immunosuppression by dexamethasone treatment was performed similar to previous protocols (8, 9). Briefly, a stock solution (12.5 mg/mL) of dexamethasone was prepared by adding 500 mg dexamethasone (Sigma-Aldrich) to 40 mL methanol (vehicle). Animals were housed in 500 mL of system water containing a final concentration of 15 mg/L dexamethasone or 0.12% methanol for 14 days. Inhibition of glucocorticoid signaling by the glucocorticoid receptor (GR) antagonist RU486 was performed using a protocol previously shown to partially phenocopy a GR mutant (10). A 20 mM solution of RU486 (mifepristone, Sigma) was prepared by dissolving 100 mg RU486 into 11.5 mL methanol (vehicle). Animals were housed in 500 mL of system water containing a 1.25 μm solution of RU486 or 0.12% methanol for 7 days. Solutions containing dexamethasone, RU486, or vehicle were changed daily and animals were fed daily with flake food approximately 2 hours prior to solution changes. At the end of the treatment, animals were euthanized and eyes were removed by enucleation and processed for immunohistochemistry.

### Notch inhibtion

Adult wild-type or *cep290* mutants were injected intraperitoneally with 20-25 μL of 1 mM RO4929097 (2,2-dimethyl-*N*-((S)-6-oxo-6,7-dihydro-5H-dibenzo[b,d]azepin-7-yl)-N’-(2,2,3,3,3-pentafluoro-propyl)-malonamide) or 1% DMSO in PBS (vehicle) using a 30-gauge needle. Animals were injected every 12 hr for 4 days and allowed to recover for 24 hr prior to euthanasia.

### RNAseq analysis

Total RNA was isolated from adult zebrafish between the age of 6-7 mpf. Animals were dark-adapted for 16-18 hours and euthanized in the dark before being moved to room light for enucleation and retina dissection Both retinas from each fish were pooled and four animals per genotype were used as biological replicates (n = 4). RNA was extracted using TRIzol with chloroform and isopropanol used to separate aqueous phases. Glycoblue (ThermoFisher) was added prior to precipitation to enhance visibility of the pellets. Nucleic acid pellets were resuspended in nuclease-free water and treated with TURBO DNase (ThermoFisher) for 15 minutes. RNA was subsequently precipitated with LiCl, washed with 70% ethanol and resuspended in nuclease-free water. RNA concentration and purity was analyzed by Qubit Fluorometer (Invitrogen) and Agilent Bioanalyzer 2100. All samples had RNA integrity number (RIN) values greater than 8.5. A total of 500 ng of each sample were submitted for Illumina RNA-seq. All library preparations, quality control steps, and next-generation sequencing was performed by the Lerner Research Institute (LRI) Genomics Core at the Cleveland Clinic. Sequencing was done on an Illumina NovaSeq 6000 with 150 bp pair-end read sequencing runs.

### scRNAseq

6 month old Cep290 and heterozygote sibling fish were utilized for all scRNAseq experiments. Whole retinas from 5 fish were dissected and dissociated as previously described (Hoang et al., 2020). Briefly, retinas were placed in Leibowitz media and incubated with hyaluronidase. Retinas were then incubated in Papain solution at 28°C with vigorous shaking for 30 minutes. Dissociated cells were then centrifuged at 1500g for 10 minutes, and resuspended in PBS containing Leupeptin and DNase. Dissociated cells were then fixed using the SplitBio single nuclei fixation kit, and samples processed using the SplitBio single cell whole transcriptome kit per the manufacturer’s instructions. Libraries were then sequenced by following the manufacturer instruction (Read1: 76, i7 Index: 6, Read2: 86) to obtain ~25,000 reads/cell. Sequencing files were processed using the SplitBio pipeline with reads mapped to the zebrafish genome GRCz11. Further processing and analysis was performed in Seurat 4.0 (Hao et al., 2021). Cells with fewer than 200 UMIs, greater than 2500 UMIs, or greater than 5% mitochondrial RNA were excluded from analysis. Known markers were used to identify retinal cell types. For differential expression analysis genes must be expressed in at least 10% of the cells within a group, have a log fold change of 0.25 or greater, and an adjusted p-value <0.05 to be considered significant.

### qRT-PCR

A total of 500 ng of purified RNA remaining from RNAseq experiments was used for reverse transcription and qPCR. Reverse transcription and qPCR were done per manufacturer’s instructions using the Bio-Rad iScript cDNA synthesis kit and the Bio-Rad SsoFast EvaGreen Supermix kit, respectively. Reactions were run on a Bio-Rad CFX96 Touch Real-Time PCR detection system and analyzed using CFX Manager. Primer sequences are listed in Table S2. Four biological replicates were prepared for each sample and three technical replicates were performed on each biological sample. Fold changes were calculated by the ΔΔC(t) method, with 18S rRNA used for normalization.

### Statistics and Data Analysis

All data was analyzed and graphed using GraphPad Prism (v8). Data sets were first tested for normal distribution using Prism. Normally distributed data sets were subsequently analyzed by Student’s t tests or one-way ANOVA with Dunnet T3 correction for multiple comparisons. Where necessary, non-parametric Kruskal-Wallace tests were used. The statistical tests and *p* values for each experiment are provided in the figure legends, with *p* values < 0.05 being considered significant.

---

**Table S6.**
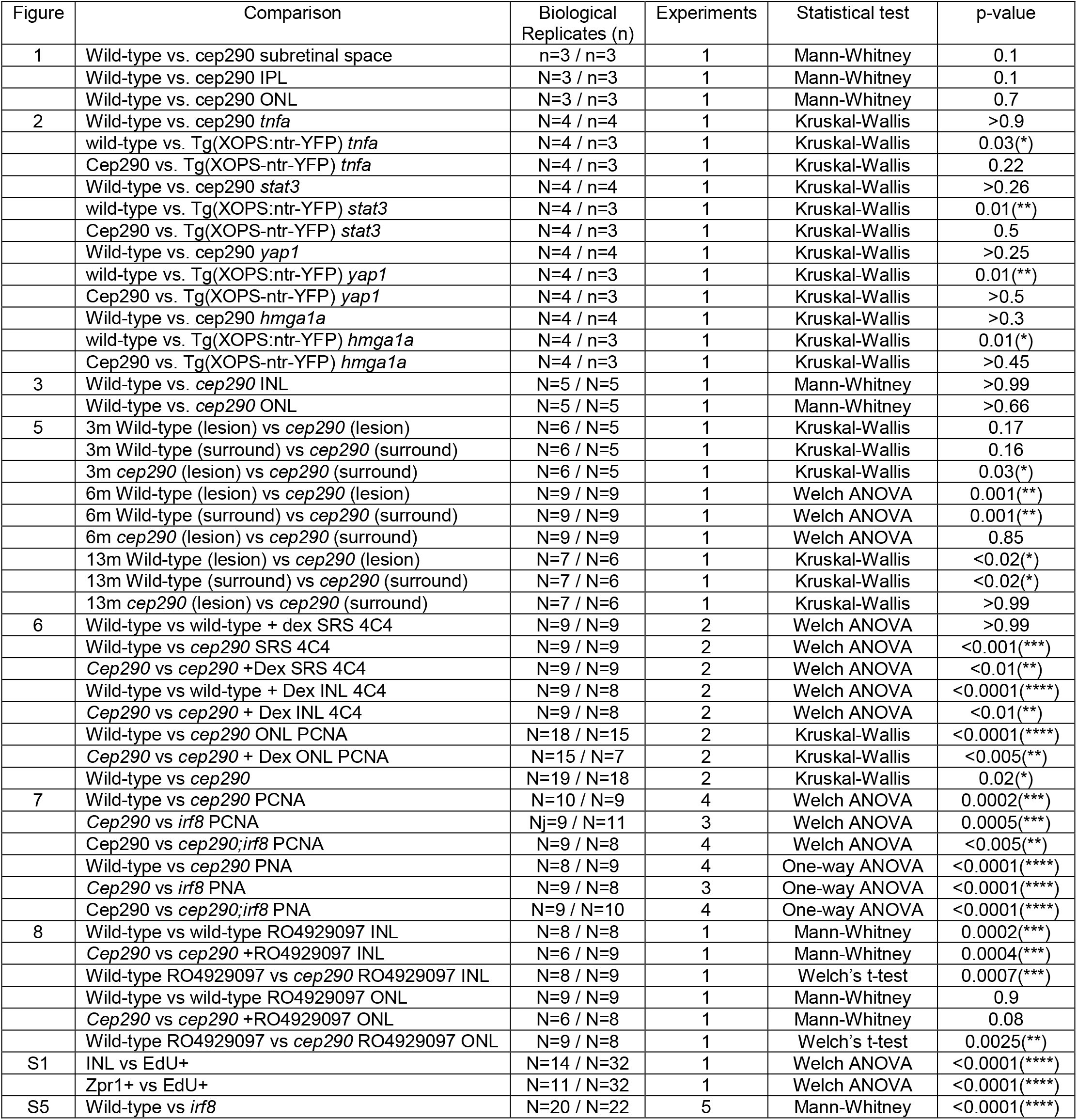
Statistical information

## Acknowledgements

This work was supported by NIH grants R01-EY017037 and R01-EY030574 to B.D.P., U01-EY027267 to D.R.H. and S.B., a Knights Templar Eye Foundation Career Initiation Grant to J.F., and a Doris and Jules Stein Professorship Award from Research to Prevent Blindness B.D.P. Additional support was provided by an NIH P30 core grant (P30-EY025585), a Foundation Fighting Blindness (FFB) Center Grant, and an Unrestricted Award from Research to Prevent Blindness to the Cole Eye Institute. The LRI Genomics Core is supported by an NIH P30 core grant (P30-CA043703). We thank Drs. Peter Hitchcock, Daniel Goldman, Jim Fadool, Ryota Matsuoka, and William Talbot for providing fish lines and reagents. We also thank the animal care staff of the Biological Resources Unit at the LRI.

## Competing Interests

**Figure S1.**
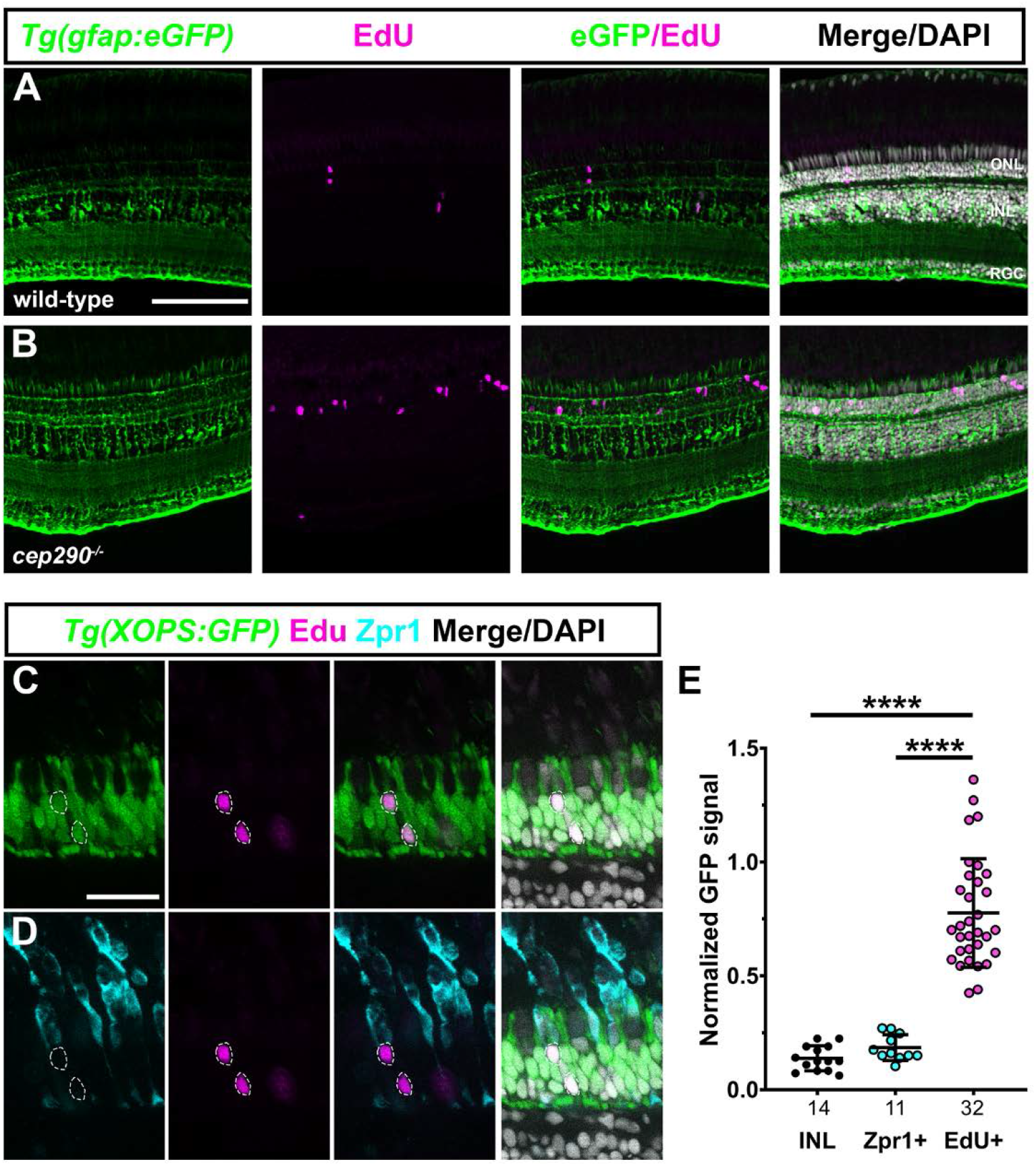
Cell proliferation occurs by rod precursors in *cep290* mutants. (A, B) EdU labeling (magenta) in retinas from 6 mpf wild-type and *cep290* mutants on the transgenic reporter line *Tg(gfap:eGFP)^mi2002^* to label Müller glia (green). Almost all EdU+ cells in *cep290* mutants were located in the outer nuclear layer (ONL). (C, D) Single optical slices showing EdU labeling (magenta) and Zpr1 immunoreactivity (cyan) in retinas from 6 mpf *cep290* mutants on the transgenic reporter line *Tg(XOPS:eGFP)*fl1, which labels rod photoreceptors (green). EdU+ cells (dashed circles) colocalized with rod nuclei but not in Zpr1+ cells. (E) The normalized GFP fluorescence signal from the *Tg(XOPS:eGFP)fl1* transgene was quantified in cells located the inner nuclear layer (INL), in Zpr1+ cells, and EdU+ cells. N-values are indicated on the graph. GFP signal was significantly higher in EdU+ cells than in Zpr1+ cells or in cells in the INL. Data are plotted as means ± SD and p-values were generated by Welch’s ANOVA test with Dunnett’s T3 multiple comparisons test. \*\*\*\**p<0.0001*. ONL = outer nuclear layer; INL = inner nuclear layer; RGC = retinal ganglion cell layer. Scale bars, 100 μm (A, B); 25 μm (C, D).

**Figure S2.**
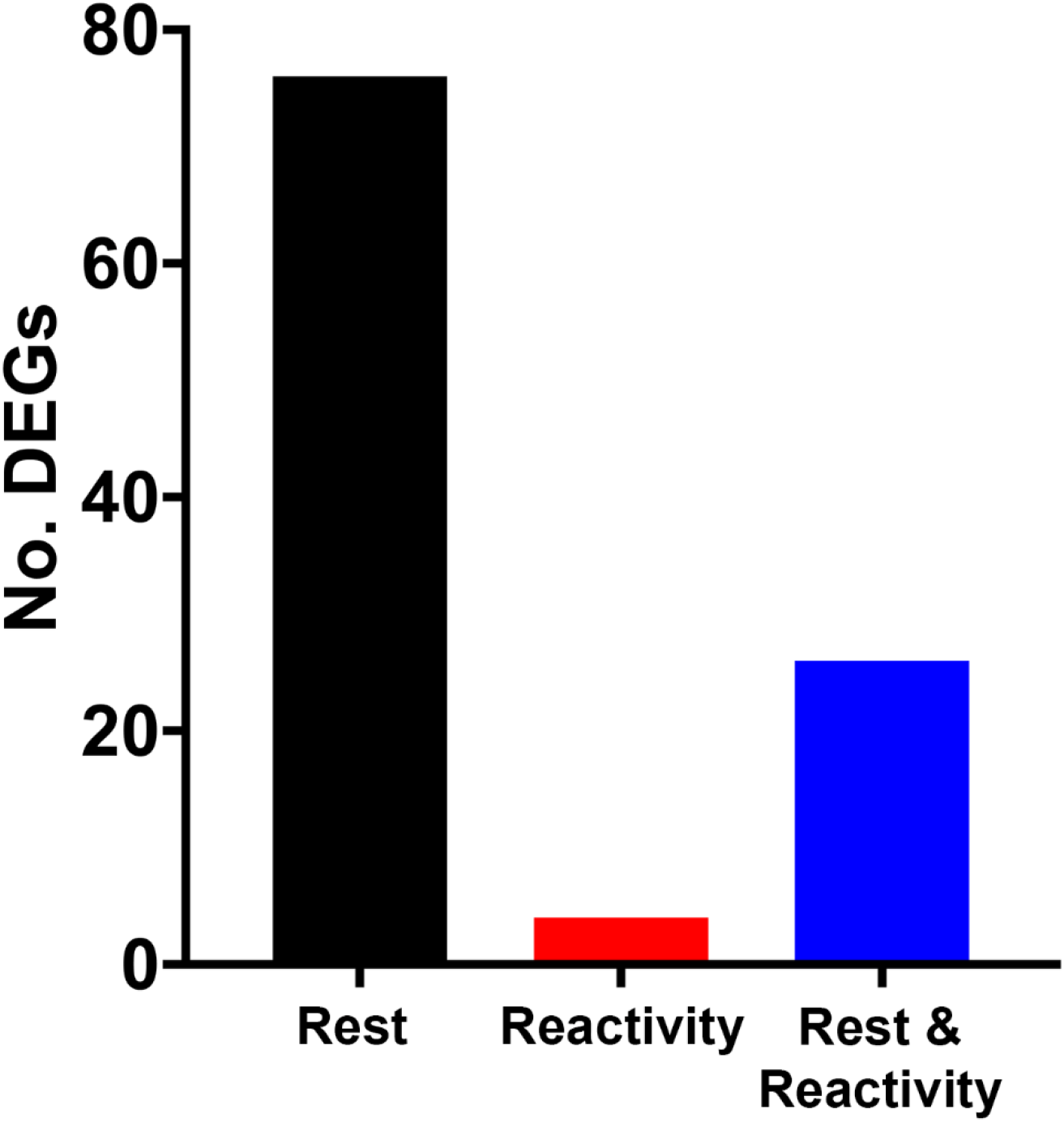
Single cell RNA-sequencing of *cep290* mutants and heterozygote siblings. Plot showing the number of differentially expressed genes associated with each pseudotime state.

**Figure S3.**
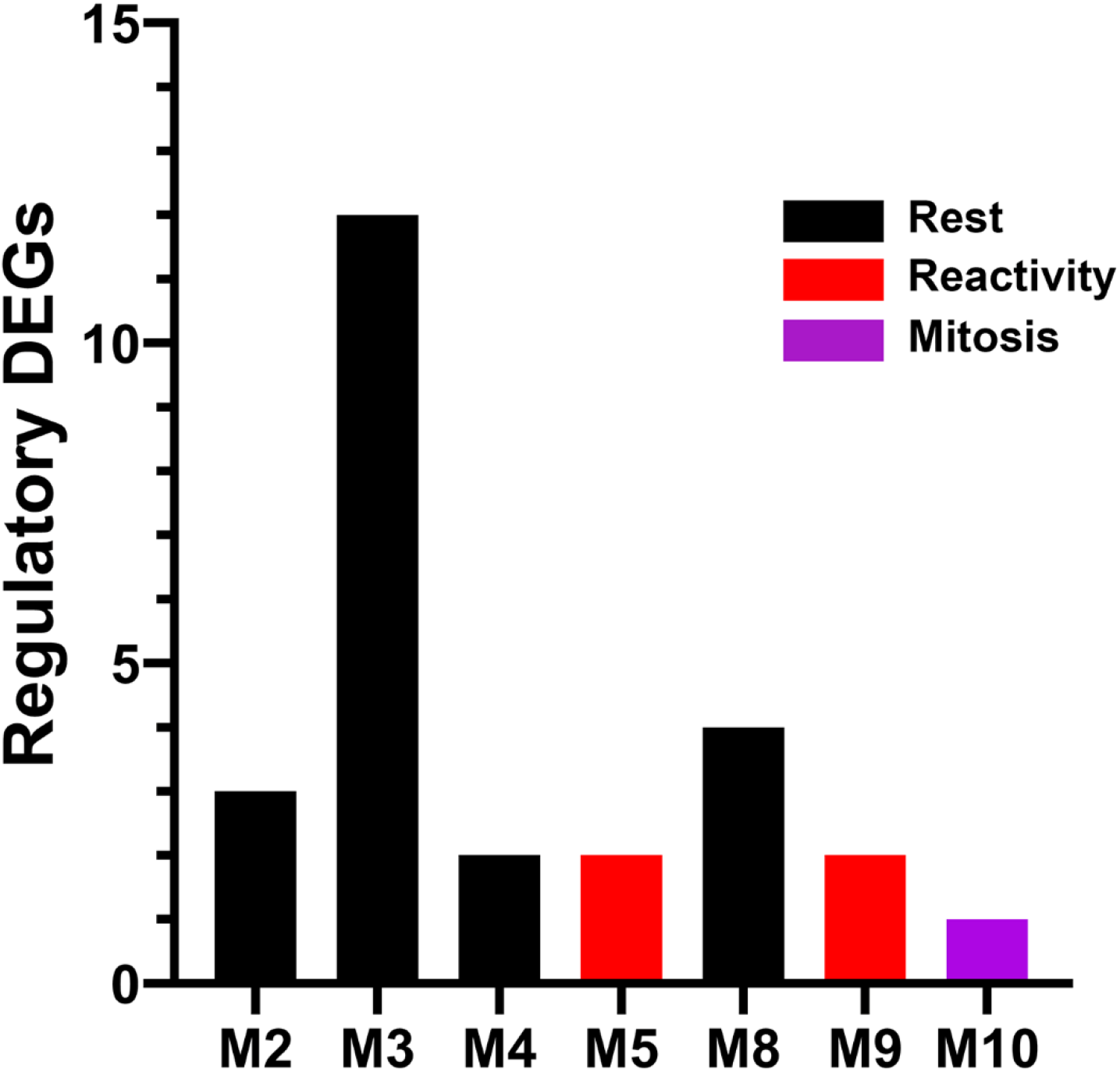
Single cell RNA-sequencing of *cep290* mutants and heterozygote siblings. (A) Plot showing the number of differentially expressed genes belonging to each regulatory module. (B) Network showing potential interaction between differentially expressed genes, all interactions show positive correlation in expression. Colors represent regulatory module.

**Figure S4.**
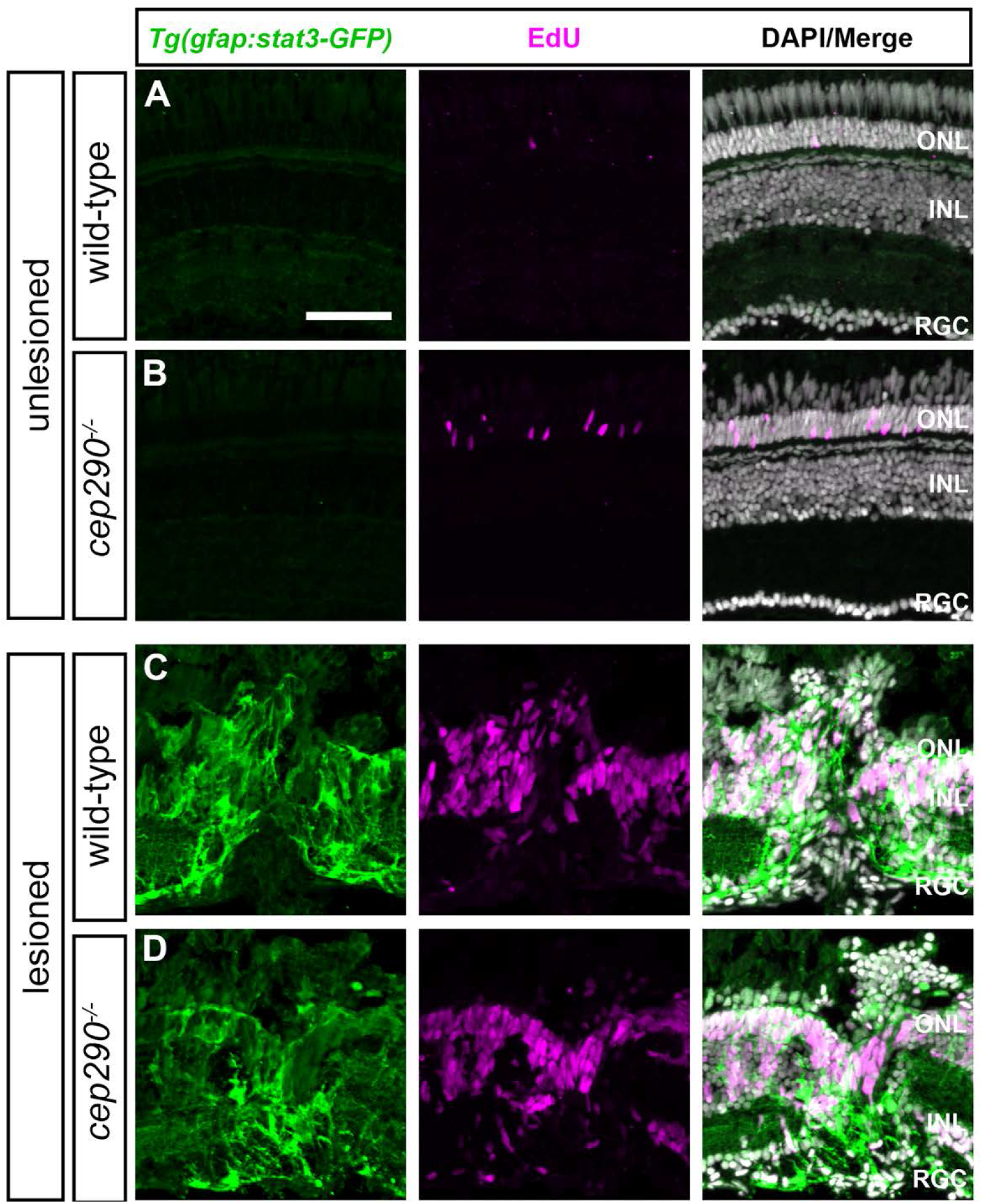
Injury is required for Stat3 activation in *cep290* mutants. Retinal cryosections of uninjured (A) wild-type and (B) *cep290* mutants on the *Tg(gfap:stat3-GFP)* background at 6 mpf were immunolabeled with anti-GFP antibodies (green) to label Stat3-GFP and processed for EdU labeling (magenta) to identify proliferating cells. In the absence of injury, Stat3-GFP expression was undetectable and the only proliferating cells were observed in the ONL. (C, D) At 4 days following mechanical injury (e.g. unilateral needle poke), Stat3-GFP expression and cell proliferation increased in both wild-type and *cep290* mutants at the site of injury. ONL = outer nuclear layer; INL = inner nuclear layer, RGC = retinal ganglion cell layer. Scale bar, 50 μm.

**Figure S5.**
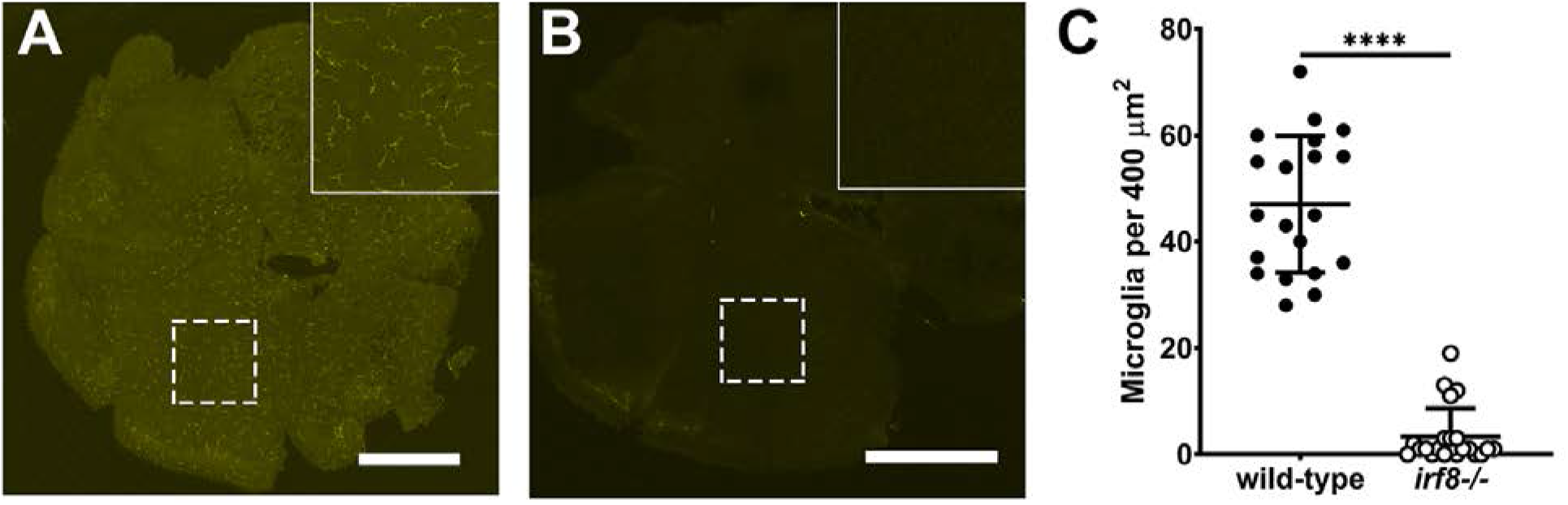
Mutation of *irf8* reduces the number of microglia/macrophages. (A-B) Confocal microscopy of flat-mounted retinas from wild-type and *irf8* mutant adults (6 mpf) immunostained with 4C4 (yellow) to label microglia/macrophage. (Inset) Higher magnification of boxed region illustrates the ramified morphology of quiescent microglia/macrophage in wild-type retinas and the significant reduction of microglia/macrophage in *irf8* mutants. (C) Quantification of 4C4+ cell density in wild-type (n = 20, filled circles) and *cep290* mutants (n = 22, open circles) found a significant decrease in microglia/macrophage. Each data point represents quantification from one complete 400 x 400 micron region of retina with 5 retinas per genotype analyzed. *irf8^-/-^* = 3.36 ± 1.13 (n = 22); wild-type = 47.05 ± 2.88 (n = 20); mean ± s.e.m. Statistical significance was determined by Mann-Whitney test (\*\*\*\**p* < 0.0001).

## Notes

https://dataview.ncbi.nlm.nih.gov/object/PRJNA787536?reviewer=ar1eg2ofco89dul9abk3i82d01

### Competing Interest Statement

The authors have declared no competing interest.

